# Standardization and Harmonization of Distributed Multi-National Proteotype Analysis supporting Precision Medicine Studies

**DOI:** 10.1101/2020.03.12.988089

**Authors:** Yue Xuan, Nicholas W. Bateman, Sebastien Gallien, Sandra Goetze, Yue Zhou, Pedro Navarro, Mo Hu, Niyati Parikh, Brian L. Hood, Kelly A. Conrads, Christina Loosse, Reta Birhanu Kitata, Sander R. Piersma, Davide Chiasserini, Hongwen Zhu, Guixue Hou, Muhammad Tahir, Andrew Macklin, Amanda Khoo, Xiuxuan Sun, Ben Crossett, Albert Sickmann, Yu-Ju Chen, Connie R. Jimenez, Hu Zhou, Siqi Liu, Martin R. Larsen, Thomas Kislinger, Zhinan Chen, Benjamin L. Parker, Stuart J. Cordwell, Bernd Wollscheid, Thomas P. Conrads

## Abstract

Cancer has no borders: Generation and analysis of molecular data across multiple centers worldwide is necessary to gain statistically significant clinical insights for the benefit of patients. Here we conceived and standardized a proteotype data generation and analysis workflow enabling distributed data generation and evaluated the quantitative data generated across laboratories of the international Cancer Moonshot consortium. Using harmonized mass spectrometry (MS) instrument platforms and standardized data acquisition procedures, we demonstrated robust, sensitive, and reproducible data generation across eleven sites in nine countries on seven consecutive days in a 24/7 operation mode. The data presented from the high-resolution MS1-based quantitative data-independent acquisition (HRMS1-DIA) workflow shows that coordinated proteotype data acquisition is feasible from clinical specimens using such standardized strategies. This work paves the way for the distributed multi-omic digitization of large clinical specimen cohorts across multiple sites as a prerequisite for turning molecular precision medicine into reality.

## Introduction

Many precision medicine projects have emerged following the launch of the Cancer Moonshot initiative, which aims to accelerate cancer research and to provide the right drugs to each patient, while also improving the prevention of cancer and its detection at an early stage. Genomic applications have undoubtedly been the major driving force for current precision medicine approaches, especially in the field of oncology, as genomic studies have revealed important and targetable cancer driver genes and mutations. However, the genotype of a patient alone is often not sufficient to support clinical decision making ^1–3^. Additional data types are necessary to bridge the gap in predicting (clinical) phenotype from genotype. Apart from clinical, lifestyle, and mobile health data, other molecular data types either alone or in combination with other available health data can support better patient diagnosis and in turn better treatment decisions.

Information about the proteotype, defined as the actual state of the proteome, the identities of proteins/proteoforms, their quantities and organization in time and space is the so-called proteogenomic data. Proteogenomics has emerged as a promising approach to advance basic, translational, and clinical research^4^. This prompted assembly of several networks and consortiums such as the Applied Proteogenomics Organizational Learning and Outcomes network (APOLLO) and the International Cancer Proteogenome Consortium (ICPC), which aim to demonstrate the critical role of proteogenomics in precision medicine and, ultimately, to incorporate proteogenomic-derived insights into patient care^5–7^. These initiatives rely on collaboration between various centers (*e.g.*, ICPC involves laboratories in multiple countries) and data sharing, which can uniquely provide information at population scale, representative of global patient diversity. Distributed data generation has been achieved in the field of genomics but not yet in the field of proteogenomics. There is thus a strong need for analytical strategies based on mass spectrometry (MS) that support proteotype analysis to deliver reliable and reproducible quantitative data that can be assembled and evaluated in a consistent and harmonized fashion. Such a capability has been demonstrated for MS-based proteotyping applications using targeted data acquisition methods and internal standards^8^, typically in clinical and late-stage translational research. However, development and assessment of standardized proteotyping workflows with acceptable quantitative performance at early stages of translational research remains a largely unmet need.

The proteome (and its many actual proteotypes at the time of measurements), is enormously complex; it has recently been estimated to include over six million proteoforms^9^. Discovery-driven clinical proteomic workflows have focused on improving coverage and reproducible quantitation of the proteome. Historic discovery methods such as label-free and isobaric-tag approaches provide both peptide identification information and relative measures of peptide abundance^10–13^. Discovery methods employing data-dependent acquisition (DDA) suffer from variable quantitative performance across samples, however^10, 11^. This led to the development of methods focused on improved stability and quantitative reproducibility such as the data-independent acquisition (DIA) techniques^14–18^. DIA-based strategies enable the unbiased measurements of peptide precursor ions (*i.e.*, MS1 spectra) as well as peptide fragment ions (*i.e.*, MS2 spectra) and leverage high-resolution and high-mass-accuracy mass spectrometry technologies to generate accurate and reproducible peptide measurements that maximize proteome quantitation. DIA-based strategies focusing on quantitation of intact peptide ion abundances extracted from retention-time aligned MS1 spectra, including accurate mass time-tag, the hybrid data acquisition and processing strategy pSMART, and hyper-reaction monitoring techniques, have demonstrated excellent analytical reproducibility^14, 19^ and quantitative accuracy^17, 20^. As these techniques afford reliable peptide quantitation and increased proteome coverage compared to DDA techniques, they are well positioned to support high-throughput clinical proteomic analyses for precision medicine applications, workflows often challenged by limited amounts of input material and large numbers of samples (hundreds or thousands) per study cohort.

To achieve reproducible and stable quantitative data sets and to facilitate harmonized implementation, standardization of DIA methods will be necessary. Toward this goal, a recent publication described the application of DIA methods to establish “digital proteome maps” of human tissue samples with the goal of creating prospective, digital proteome biobanks of clinical biospecimens supporting real-time and retrospective data analyses^21^. Moreover, optimized synthetic^22^ and internal^23^ peptide standards have been developed to facilitate peptide retention-time alignment procedures and support facile comparison of DIA datasets generated at different analytical sites. Further, efforts to benchmark software platforms^24^ and statistical methods^25^ for DIA data analysis have been described as has the generation of comprehensive peptide spectral libraries from diverse human tissues and cell lines for DIA analyses of human samples to support standardized DIA data processing^26^. Recently, performance benchmarks for DIA data acquisition, specifically the application of the so-called SWATH DIA-MS approach, were used in analyses of a complex cell line standard in 11 laboratories in nine countries^27^. This study revealed consistent quantitation of more than 4000 proteins from HEK293 cells across all laboratories; the analyses also involved quantification of a panel of internal peptide standards over several orders of magnitude in concentration. It demonstrates that DIA-MS data acquisition is a reproducible method for large scale protein quantification, while in our multi-national study, our efforts were focused on establishing a comprehensive label-free quantitation DIA-MS workflow with a much higher throughput to address the needs of large cohort clinical sample profiling. A quality control system was developed to monitor the entire workflow performance, determine the instrument performance gap, and guide to a promptly troubleshooting when necessary, ensuring high data quality and secure the throughput necessary for a large cohort study. Label-free quantitation performance was evaluated with a well-established label-free quantitation sample set at each laboratory through the entire study, in order to provide a benchmark standard using the standardized HRMS1-DIA workflow across different labs and days, enabling the data sets acquired in a longitudinal mode and/or at different sites can be compared and normalized during big data analysis.

To continue to expand the implementation of DIA workflows and integrate them into routine clinical sample analyses, the incorporation of benchmarked standards and standardized quality control routines are necessary to maximize the accessibility of downstream data and empower team-driven science initiatives. Here we report on the performance of a quality control (QC) and biological sample data acquisition schema analyzed using a streamlined QC-benchmarked HRMS1-DIA workflow implemented in a continuous operational mode for seven consecutive days in eleven labs in nine countries followed by centralized data processing. Controlled samples^24^ included *E. coli*, yeast and human cell line peptide digests combined at fixed ratios and were used to mimic biological samples and provide proof-of-concept feasibility and performance of this streamlined workflow. Experiments were then extended to actual biological samples from well-defined ovarian cancer histotypes, namely high-grade serous and clear cell ovarian cancers, to further demonstrate the utility of the standardized HRMS1-DIA strategy for routine clinical proteotype analyses.

## Results

### Implementation of a QC-Benchmarked, HRMS1-DIA Mass Spectrometry Workflow

This report details the analytical performance and reproducibility of a standardized, QC-benchmarked HRMS1-DIA workflow intended to achieve quantitative proteotype analysis studies of large cohorts across several centers and in turn to support precision medicine projects. To address the needs of a robust and high-throughput workflow compatible with large-cohort studies, a 60-minute capillary flow LC gradient using 1.2 µl/min analytical flow rate was applied in all the analyses. Primary data acquisition was performed on either the Easy-nLC 1200 or the Ultimate 3000 RSLC nano liquid chromatography system coupled to Q Exactive HF mass spectrometers (Thermo Fisher).

The HRMS1-DIA method used involves multiple MS1 scans interspersed with 18 DIA MS/MS scans per duty cycle (in total 54 DIA MS/MS scans) (**Figure 1A**). Quantification was based on precursor ion signals measured through high-resolution full MS scans with 120k resolution setting; the MS2 information was utilized for peptide identification only. The MS1 scan repetition rate was set independently of the MS2 cycle time such that a sufficient number of data points were acquired over peptide chromatographic elution profiles for a proper determination of peptide peak areas and therefore their precise quantification. The overall MS2 cycle length was maximized under the condition that each parent ion was sampled approximately three times within the duration of a typical chromatographic peak. This allowed the precursor isolation windows to be minimized for the DIA MS/MS scans (*i.e.*, 15 *m/z* units in this study). This moderate precursor isolation window width directly enhances the selectivity and confidence of the MS/MS-based peptide identification. In order to systematically evaluate the reproducibility of the QC-benchmarked HRMS1-DIA workflow, spectral libraries were centrally built from the DDA analysis of high pH reverse phase fractions. Spectronaut software (Biognosys) was applied for both individual onsite data analysis and central data analysis with the centrally prepared spectral libraries.

**Figure 1.**
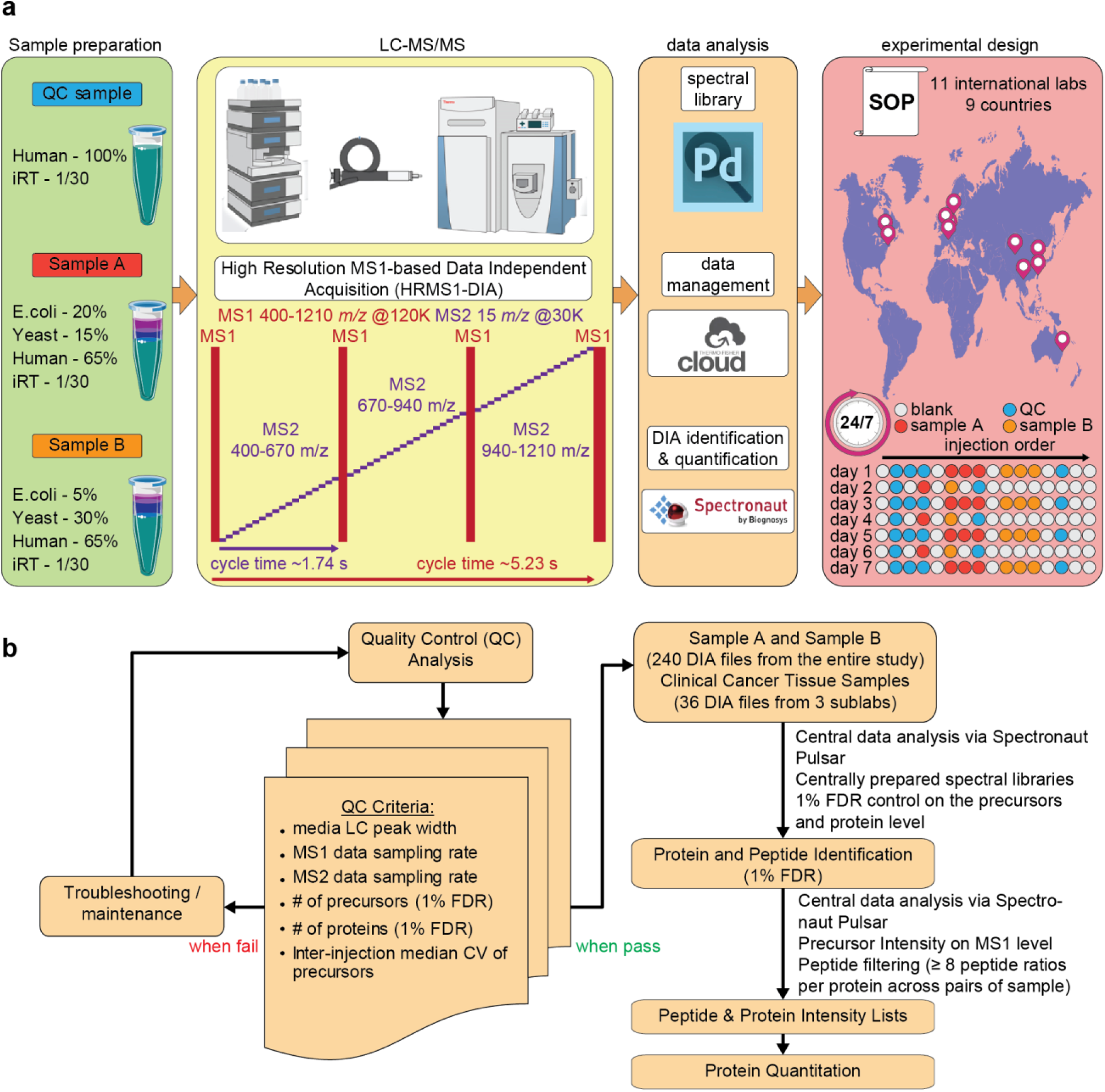
QC standard and controlled sample data acquisition and analysis using a streamlined HRMS1-DIA workflow: **A**: QC Sample is HeLa lysates. Sample A and B are mixtures of HeLa, yeast, and *E. coli* lysates. The samples were analyzed by capillary LC-HRMS1-DIA on a Q Exactive HF system at eleven sites in a 24/7 mode for 7 consecutive days. On Days 1, 3, 5, and 7, all samples were run in three technical replicates; on Days 2, 4, and 6, Samples A and B were each run once, and the QC standard was run once before and after the Sample A and B. The rest of the time, the instruments were running blank injections. **B**: Quality control criteria were based on the evaluation of the performance of the capillary LC-HRMS1-DIA workflow before the 11 labs kicked off the study. During the study, a QC standard was analyzed in three technical replicates on Days 1, 3, 5, and 7. If the QC criteria were met, Sample A and Sample B data sets acquired on the same day were analyzed. If the QC standard analysis did not pass QC criteria, either instrument setup maintenance or troubleshooting were undertaken. In total, 240 DIA files from both Sample A and Sample B were centrally analyzed via Spectronaut v11 with centrally generated spectral libraries. A criterion of 1% FDR was applied for identification at precursor and protein levels. The intensity of each identified peptide was exported to an .xls file, which was further processed via an R-script with the “peptide-to-protein rollup pairwise ratio” quantification strategy (See “Methods” section).

The workflow is a two-step procedure. First, the performance of the LC-MS platform operated with the HRMS1-DIA acquisition method was assessed in order to detect “drifts” from the predefined performance baseline and to allow corrective measures to be taken if needed. A QC standard was analyzed using rigorous metrics to support system suitability testing; the QC standard was a commercially available peptide digest derived from the HeLa human cervical cancer cell line. Second, following successful platform qualification, the actual quantitative measurements of samples of interest were performed using the same acquisition method.

The baseline for the system suitability test was generated from the analyses of the QC standard performed by four reference laboratories in continuous operation mode over several days with LC-MS platforms operating at different levels of performance (data not shown). The collection of such data enabled the establishment of reference metrics and associated acceptance criteria for platform qualification from three replicate analyses of the QC standard. Reference metrics included median LC peak width, MS1 and MS2 sampling rates, total precursor ions and protein groups identified, and inter-injection median CV on precursor ion signals (**Table 1**). These metrics provided reliable read-outs and real-time monitoring of platform status, covering both chromatographic and mass spectrometric attributes. These QC acceptance criteria were also applied to diagnose possible issues such as need for maintenance in order to maintain high-throughput for the vast sample sizes in the studies.

**Table 1.** Reference metrics and associated acceptance criteria from the system suitability QC tests

|  |  |  |  |  |  |
| --- | --- | --- | --- | --- | --- |
| Median LC Peak Width (s) | MS1 Data Sampling Rate | MS2 Data Sampling Rate | Precursor IDs (1% FDR) | Protein IDs (1% FDR) | Inter-injection median CV of precursors |
| $17 \pm 3$ | 8-11 | 3-4 | $> 48000$ | $> 5000$ | $\leq 15\%$ |

### Performance Evaluation of QC-benchmarked HRMS1-DIA Workflow Applied to Controlled Samples

The QC-benchmarked HRMS1-DIA was applied to the determination of abundances of peptide digest mixtures of defined composition, referred to as controlled samples, by each of the 11 research laboratories in a 24/7 operation mode for seven consecutive days (**Figure 1A**). The controlled samples were mixtures of digests prepared from diverse organisms: Sample A was 65% HeLa, 15% yeast, and 20% *E. coli* and Sample B was 65% HeLa, 30% yeast, and 5% *E. coli*. Each laboratory followed standard operating procedures (see standard operating procedures in Supplementary Protocols). Each day, system suitability tests of the QC standard were performed, and no action was taken as long as QC acceptance criteria were satisfied (**Figure 1B**). On Days 1, 3, 5, and 7, the QC standard was analyzed in triplicate and then the control samples were analyzed in triplicate. At the end of each day, the QC standard was run. On Days 2, 4, and 6, the QC standard was run once at the beginning and end of the day and the controlled samples were also analyzed once (**Figure 1B**).

At each laboratory, the files of data obtained on the QC standard were processed daily with Spectronaut Pulsar, enabling the extraction of QC metrics evaluated in system suitability tests (**Supplementary Table 1**). The QC files were searched against a spectral library constructed from DDA analyses of a peptide digest derived from human cell line KG1a (See “Methods” section).

The QC acceptance criteria were systematically satisfied for analyses performed by nine of the eleven laboratories, translating into the identification of 5028 to 5993 protein groups with 1% FDR (**Figure 2**). One laboratory (Lab 10) faced major challenges, mainly resulting from poor chromatographic separations, which could not be resolved under the time constraints of the study. Another participating laboratory (Lab 5) experienced some technical issues on Day 7, translating into lower overall performance; specifically, only 4423 protein groups were identified. As the median LC peak width, sample rates at both MS1 and MS2 level, and inter-injection median CV on precursor ion signals were within the established criteria, the performance issues were not related to the chromatographic separation. Further investigation revealed the need for maintenance of the HCD cell of the mass spectrometer. After necessary maintenance on Day 8, the normal operation performance was recovered on Day 9. This demonstrates that the QC analysis enables real-time monitoring of instrument status, identifies performance gaps, and provides a guide to root causes of performance issues, enabling the high-throughput performance needed for large cohort studies. Therefore, of the 11 participating laboratories, 10 were able to perform sample analyses on Days 1, 3, 5, and 7, although Lab 5 continued on Day 9 instead of Day 7. Lab 10 did not participate in controlled sample analyses.

**Figure 2:**
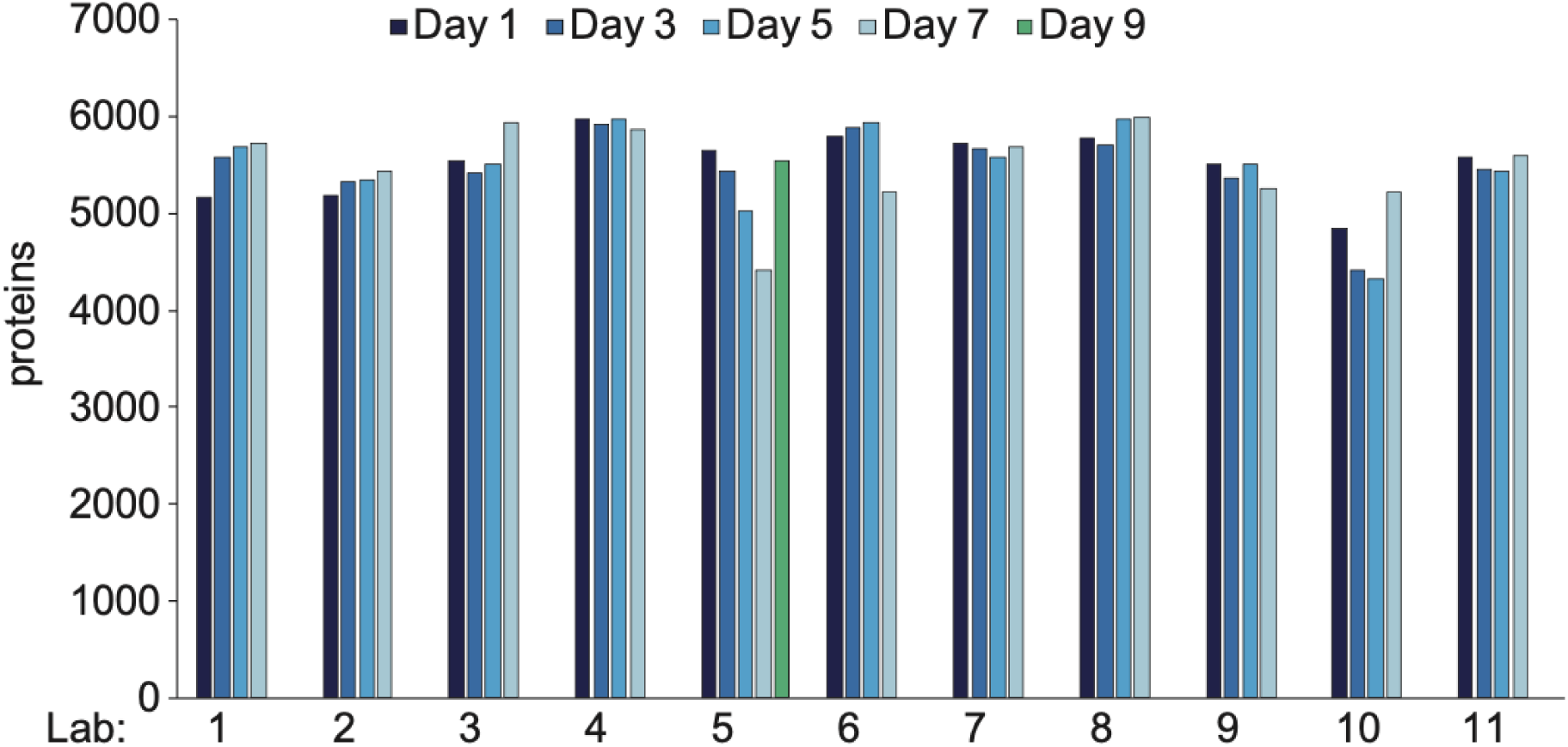
Quality Control Performance. The number of proteins identified (with a 1% FDR) from the HRMS1-DIA analyses of QC sample (2 µg of Hela digest on column) prior to controlled samples analyses at each laboratory was plotted over the 7-day evaluation period. The evaluation period was expanded to 9 days (green bar) for laboratory 5, experiencing technical issues on day 7 which were subsequently addressed.

All data generated from Samples A and B were centrally processed in Spectronaut Pulsar. For peptide/protein identifications, the human spectral library used for QC standard data processing was supplemented with similarly constructed yeast and *E. coli* libraries. Peptide precursors were quantified using mass chromatogram areas extracted from MS1 data, and protein abundance changes were determined using a strategy in which a minimum of eight pairwise peptide ratios were combined across technical replicate injections for a given protein group (**Supplementary Table 2**). A total of 240 DIA injections of the Samples A and B were acquired at 10 research sites that met QC criteria, and more than 7600 protein groups were identified with 1% FDR (**Figure 3A**). Approximately 4000 human proteins, 2000 yeast proteins, and 400 *E. coli* proteins (**Figure 3B**) were quantified across the three injections per day over the four data acquisition days.

**Figure 3.**
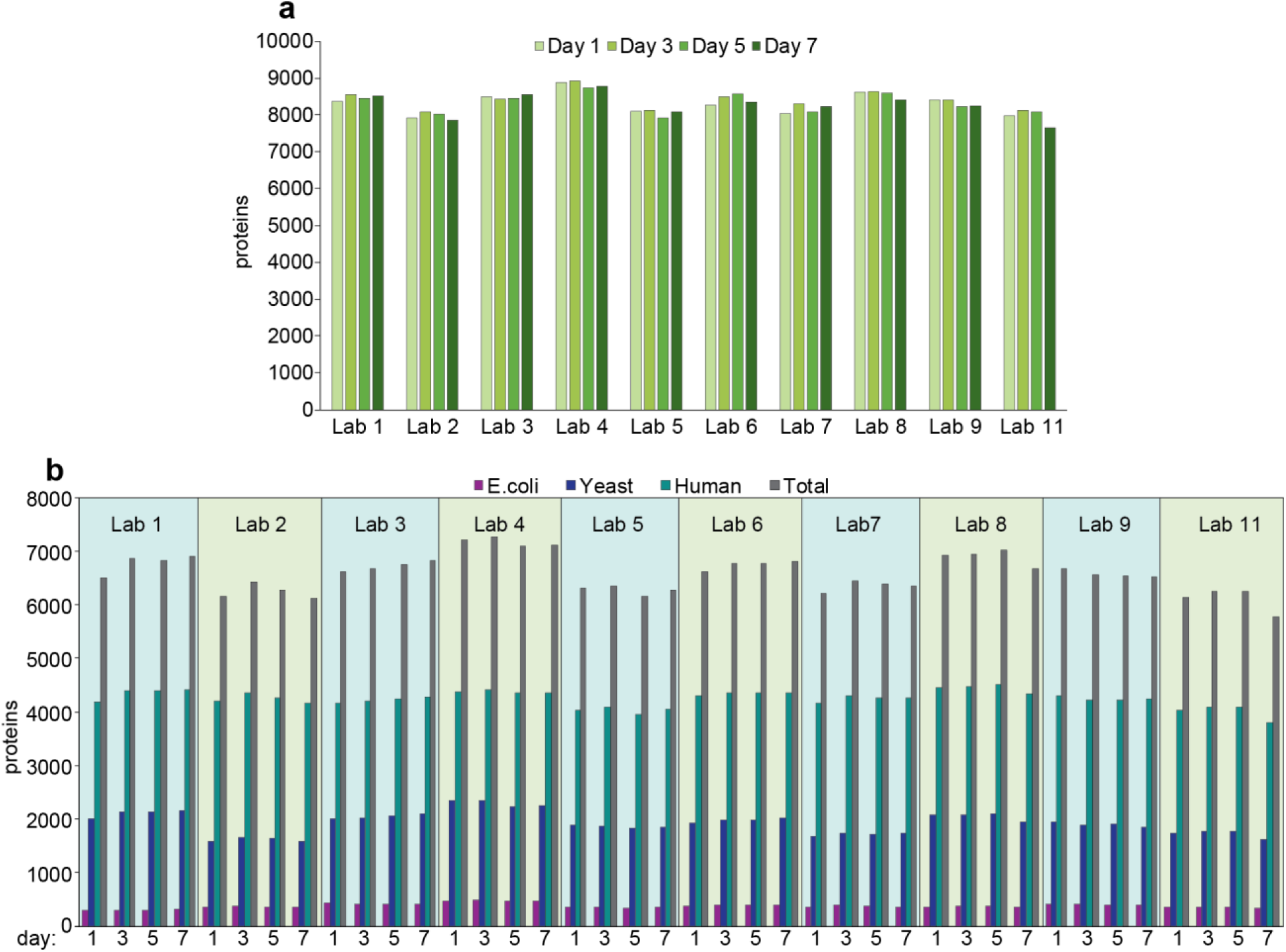
Overall performance of QC-benchmarked HRMS1-DIA workflow based on controlled samples analyses. **A**: The number of proteins identified in both samples A and B analyses (with a 1% FDR) at each laboratory was plotted over the 7-day evaluation period. **B**: The number of proteins quantified in total and for each individual organism in controlled samples (based on criteria described in “Methods” section) at each laboratory is plotted over the 7-day evaluation period. For laboratory 5, benefiting from an extended evaluation period of 9 days, due to technical issues detected and addressed during day 7, the identification (A) and quantification (B) results obtained for day 9 substitutes those of day 7.

The inter-day reproducibility was excellent at each site as more than 80% of the total number of locally quantified protein groups were quantified on each of the four data acquisition days. On average, more than 6500 protein groups were quantified with a relative standard deviation (RSD) < 4% at each site (**Figure 4A**). Furthermore, the inter-lab reproducibility was comparable with a total of 5784 proteins groups quantified across the labs, representing approximately 80% of the proteins quantified locally. Notably, 4565 of these protein groups were not only quantified by all sites but were also quantified on each acquisition day (**Figure 4B**).

**Figure 4.**
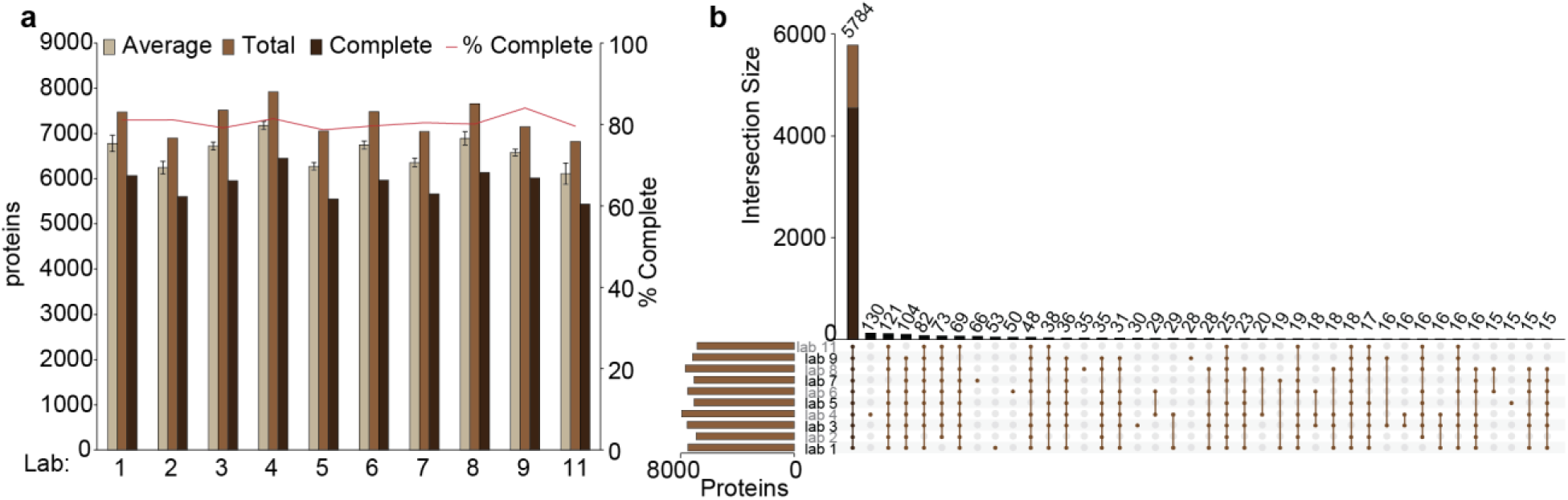
Reproducibility of quantitative proteome coverage achieved by HRMS-DIA analyses. **A**: The number of proteins quantified in controlled samples at each lab (based on criteria described in “Methods” section) in total, on average, and systematically across the 7-days evaluation period was plotted together with the estimation of the proportion of complete profile (“Complete”/”Total”). **B**: The numbers of proteins co-quantified across various combinations of laboratories were plotted (vertical brown bars). The vertical dark brown bar (4565 proteins) overlapping brown bar (5784 proteins) for the combination including all the laboratories reflects the number of proteins co-quantified by all the laboratories every evaluation day. The horizontal brown bars reflect the number of proteins quantified in total at each laboratory.

In-depth evaluation of quantitative performance relied on the experimentally determined abundance differences between controlled samples A and B. The results demonstrated high quantification accuracy compared to theoretical abundance differences as reflected by the low deviation of experimental values; the median values were typically lower than 10% for human and yeast proteins and typically lower than 20% for *E. coli* proteins (**Figure 5, upper panel**). The highest deviation from theoretical values was observed in data collected by Lab 1, which may have resulted from some inaccuracy in sample preparation. The analytical quantitative precision was excellent for the 10 participating laboratories, as illustrated by median CV values that were typically below 5% for the human and yeast proteins and below 10% for the *E. coli* proteins (**Figure 5, lower panel**). The slightly higher value obtained for the *E. coli* proteins likely resulted from the overall lower abundance of the *E. coli* proteome in the sample mixtures.

**Figure 5:**
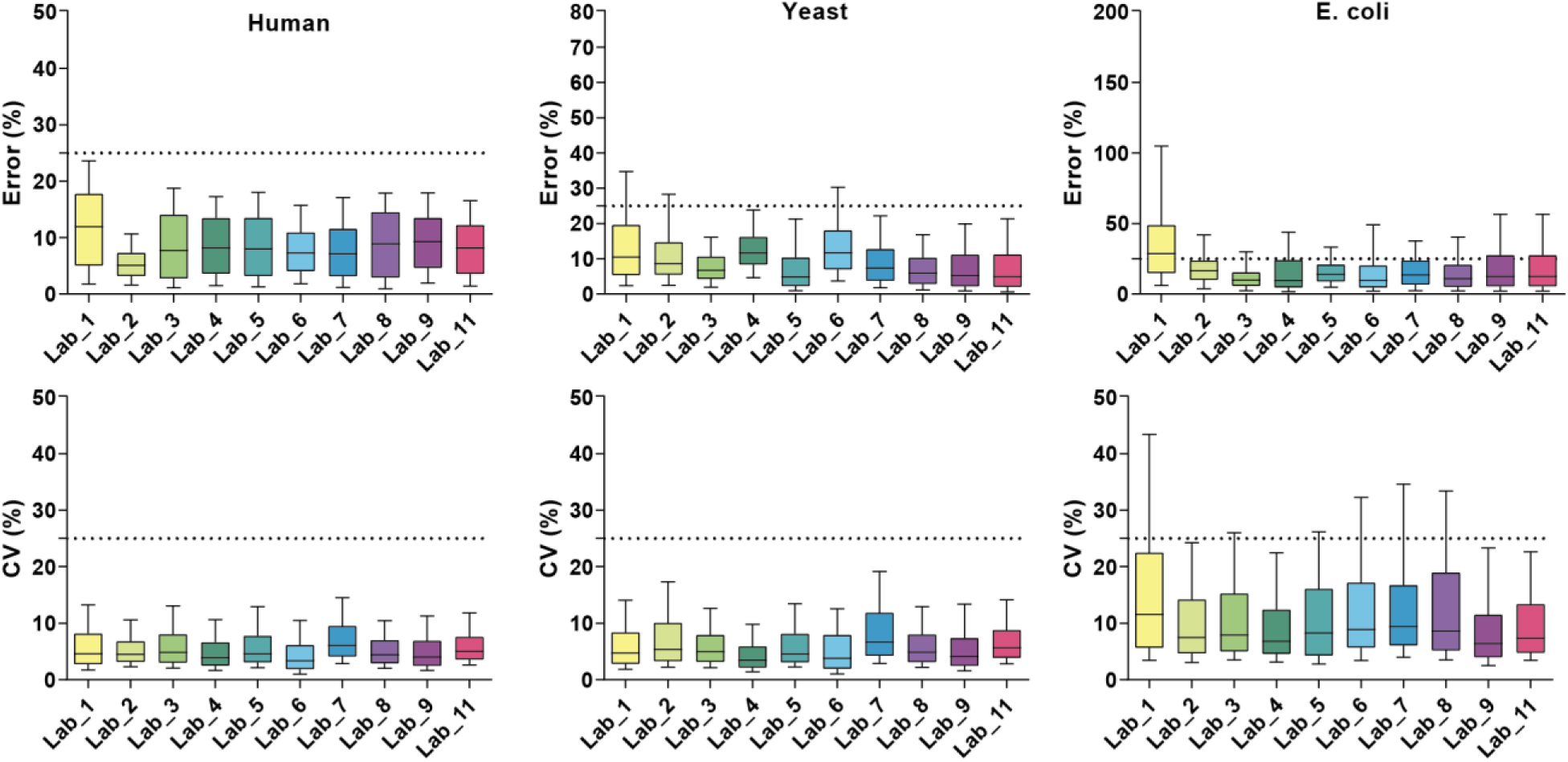
Analytical precision and accuracy of protein quantification from the analyses of controlled samples. The evaluation was based on the experimentally determined abundance changes of the systematically quantified proteins (every day by every laboratory) between controlled samples A and B. In the upper panel, the distribution of the deviations from theoretical protein abundance changes for the three organisms (error, in %) was plotted for each laboratory through boxplots with whiskers representing the 10^th^ and 90^th^ percentile. Horizontal dashed lines were added at 25% error, as a reference value. In the lower panel, the distribution of the coefficients of variations obtained on the determined protein abundance changes for the three organisms across the various evaluation days (CV, in %) was plotted for each laboratory through boxplots with whiskers representing the 10^th^ and 90^th^ percentile. Horizontal dashed lines were added at 25% CV, as a reference value.

### Analyses of Tumor Tissue Digests by QC-benchmarked HRMS1-DIA to Confirm Disease-Relevant Protein Alterations between Histotypes

This standardized methodology was also applied to the analysis of complex tissue digests prepared from ovarian cancer tumors in three of the participating laboratories. Formalin-fixed, paraffin-embedded (FFPE) tissues, derived from two high-grade serous ovarian cancer (HGSOC) and two clear-cell ovarian cancer (OCCC) tumors, were sectioned onto polynapthalate membrane slides, and tumor cell populations were harvested by laser microdissection (**Figure 6A**). Enriched epithelial cancer cell populations of interest were digested with trypsin using a pressure cycle technology workflow, and resulting peptides were analyzed using the QC-benchmarked, HRMS1-DIA method described herein in three technical replicates at each of three independent analytical laboratories. Data were centrally processed, and protein-level abundances were determined using eight peptide ratios spanning tumor datasets by disease histotype.

**Figure 6:**
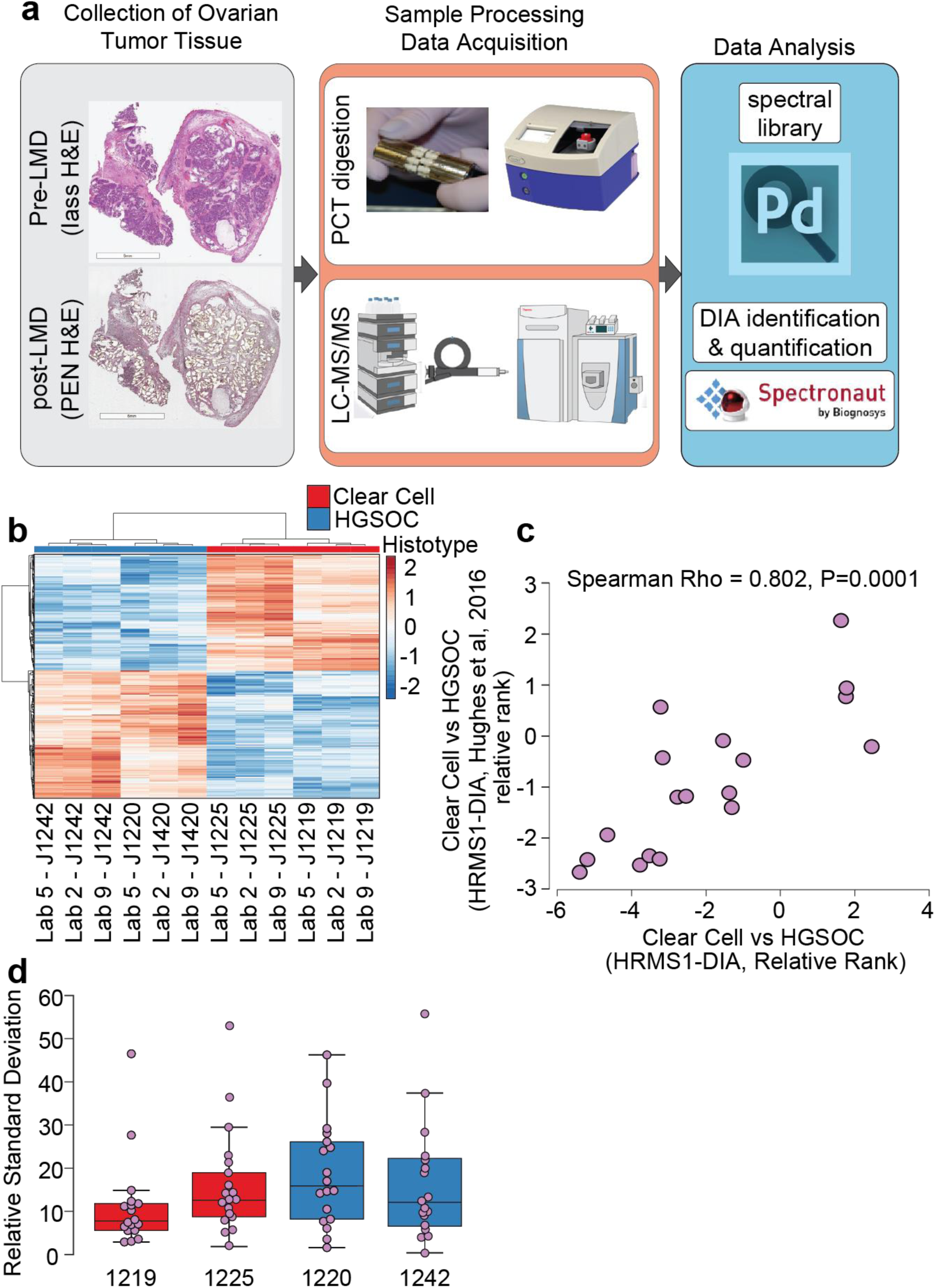
Analyses of HGSOC and OCCC samples using HRMS1-DIA at three analytical sites. **A:** Epithelial cancer cells laser microdissected from HGSOC and OCCC tumors were evaluated by three labs from three different analytical sites using the standardized protocol. Each biological sample was measured as three technical replicates. QC standards were evaluated before, between, and after cancer tissue samples. The data acquisition was performed in a 24/7 operation mode. **B**: A heatmap of reflecting unsupervised cluster analysis (Pearson correlation and average linkage) of the 394 proteins that were significantly differentially expressed (LIMMA adjusted p-value < 0.01) between HGSOC and OCCC tissues. **C**: Correlation of expression in HGSOC versus OCCC tissues of proteins from the gene signature identified by Hughes et al (insert citation here). We co-quantified 18 significant protein alterations (LIMMA p-value < 0.05) with a 112 gene signature stratifying HGSOC and OCCC ovarian cancers. Protein alterations were significantly correlated with this feature subset (Spearman = 0.802, p=0.0001). **D**: Quantitative reproducibility of 18 HGSOC and OCCC signature proteins across three analytical sites by patient tissue sample. The figure shows the relative standard deviation between protein abundances obtained for the 18 signature proteins quantified at the three laboratories.

A total of 5712 proteins were quantified across tumor samples (**Supplementary Table 3**); 394 proteins were expressed at significantly different abundances in HGSOC and OCCC tissues (**Supplementary Table 4; Figure 6B**). These differentially expressed proteins were compared with a protein signature previously reported to stratify HGSOC and OCCC^28^, and 18 co-quantified proteins were found to exhibit concordant protein abundances (R=0.802, P=0.0001) (**Figure 6C**). Quantitative reproducibility of 18 HGSOC and OCCC signature proteins across three analytical sites for each patient tissue sample revealed a mean relative standard deviation of 15.45 with a standard error of ± 1.43 (**Supplementary Table 5; Figure 6D**).

## Discussion

A standardized analytical approach to support the creation of digital proteogenomic biobanks of clinical biospecimens for prospective and retrospective data analyses has been designed. To realize this goal, standardized and quality-control-benchmarked workflows were necessary. The developed approach for the generation of defined datasets will maximize the consumption and (re)usability of downstream data sets and empower team-driven science initiatives and population-level insights between the proteome and human disease. Our international and multi-site study defined a quality control-driven, HRMS1-DIA workflow with excellent analytical reproducibility and label-free quantitative performance providing deep global proteotype profiling across diverse sample types including clinical tissue samples. The workflow resulted in the consistent and confident identification of more than 5000 proteins in the peptide digest derived from a human cell line that served as our QC-benchmark standard. Applied to the analysis of the complex mixture of digests from human, yeast, and bacterial cells, it allowed identification, on average, of more than 7600 proteins and the quantitation of more than 6500 proteins for ten of the eleven laboratories that participated.

We provided further proof of concept by applying this workflow to tryptic digests established from “real-world” FFPE cancer tissue specimens. Highly reliable protein quantitation enabled the detection of disease histotype-specific protein alterations in each of the three laboratories that took part in this exercise. The consistent depth of proteome coverage (>80% of total proteins quantified across partnering sites) and analytical performance achieved across diverse sample types and sites using a 1-hour, capillary LC-MS method operating in a 24/7 mode demonstrates that this high-throughput and robust workflow, enabling the quantification of more than 100 proteins per minutes, is ready for application to large-cohort tissue proteomic studies. This workflow defines and implements QC benchmark expectations that can be monitored in real-time during primary DIA data production to ensure that data quality standards are achieved and that can be leveraged during downstream data processing to assess and track analytical variability and bias in cohort-level data. The presented HRMS1-DIA workflow is a standardized, quantitative method that is driven by defined quality-control expectations and that exhibits stable, highly reproducible, and scalable performance to support both basic discovery proteomics research and population-scale clinical sample analyses in a high-throughput manner.

## Supporting information

Standard operating procedure - Sample preparation

Standard operating procedure - DIA Analysis with Capillary-flow UltiMate 3000 RSLCnano

Standard operating procedure - DIA Analysis with Easy nLC 1200

Standard operating procedure - DIA Analyses and Data Evaluation

QC-DIA benchmark proteins quantified by analytical site.

MultiSite proteins quantification results of controlled samples (Log2 ratio protein abundance (Sample A / Sample B).

Tumor tissue proteins quantified by analytical site.

Significantly altered/ histotype correlated protein alterations.

The relative standard deviation (RSD) of 18 HGSOC and OCCC signature proteins across three analytical sites for each patient tissue sample.

## Abbreviations

QC: Quality Control
MS: Mass spectrometry
HRMS1-DIA: High-resolution MS1-based quantitative data-independent acquisition
DIA: Data-independent acquisition
FFPE: Formalin-fixed, paraffin-embedded
DDA: Data-dependent acquisition
HGSOC: High-grade serous ovarian carcinoma
OCCC: Ovarian clear-cell carcinoma

## Methods

### QC Standard

The QC standard was a commercially available peptide digest derived from the HeLa human cervical cancer cell line, the Pierce HeLa Protein Digest Standard (Thermo Fisher Scientific), resuspended at a concentration of 1 µg/µL in HPLC-grade water containing 0.1% (v/v) formic acid and supplemented with eleven non-naturally occurring synthetic peptides from the iRT kit (Biognosys) at a ratio 1:30 v/v.

### Mixed Proteome Samples

The controlled samples A and B were prepared from the Pierce HeLa Protein Digest Standard (Thermo Fisher Scientific), the Mass Spec-Compatible Yeast Digest (Promega), and the MassPREP *E. coli* Digest Standard (Waters). Each digest was resuspended at a concentration of 1 µg/µL with HPLC-grade water containing 0.1% (v/v) formic acid. Sample A was prepared by mixing human, yeast, and *E. coli* protein digests at 65%, 15%, and 20% w/w, respectively. Sample B was prepared by mixing human, yeast, and *E. coli* protein digests at 65%, 30%, and 5% w/w, respectively. The iRT kit (Biognosys) was added to each of the controlled samples at a ratio 1:30 v/v. For analysis, 2 µl of each sample was injected. The detailed preparation instructions are described in Supplementary Protocol 1.

### Cancer Tissue Specimens

Four FFPE ovarian cancer surgical tissue specimens (two OCCC and two HGSOC) from an IRB-approved protocol were selected for laser microdissection. Thin (10 µm) tissue sections were cut using a microtome and placed on polyethylene naphthalate membrane slides. After staining with aqueous hematoxylin and eosin, laser capture microdissection (Leica LMD7) was used to harvest tumor cells from thin sections, which were collected by gravity into microcentrifuge tubes containing 45 µL of LC-MS grade water (mean tumor cell area captured = 86.6 ± 1.7 mm^2^). Tissue samples were vacuum dried and transferred into MicroTubes (Pressure BioSciences, Inc., Medford, MA) containing 100 mM TEAB/10% acetonitrile (ACN) and incubated at 99 °C for 30 min. The temperature was lowered to 50 °C, and 1 µg of porcine trypsin was added to each tube. Tubes were capped with MicroPestles (Pressure BioSciences, Inc.). Pressure-assisted digestion was performed in a 2320EXT barocycler (Pressure BioSciences, Inc.) by sequentially cycling between 45 kpsi (50 s dwell time) and atmospheric pressure (10 s dwell time) for 60 cycles at 50 °C. The peptide digests were transferred to 0.5-mL microcentrifuge tubes, vacuum dried, and purified using Pierce C_18_ Spin Columns. Resulting peptides were resuspended in 25 mM ammonium bicarbonate, pH 8.0 and the peptide concentration of each digest was determined using the bicinchoninic acid (BCA) assay (Pierce Biotechnology).

### Cell Culture and Lysis

A human cell line KG1a was maintained and propagated in RPMI 1640 medium supplemented with 10% FBS and 1% penicillin/streptomycin at 37 °C in a 5% CO_2_ atmosphere, and cells were lysed in buffer containing 4% SDS. Lysates were sonicated (Ningbo Scientz) and centrifuged at 15,000 × g for 15 min. The protein concentration of the supernatant was determined by using the BCA assay kit. Proteins (200 µg) were digested overnight with Lys-C (Wako Chemicals) and trypsin (Promega) using the filter-aided sample preparation protocol^29^. Peptides were recovered and desalted using Oasis HLB 1-cc cartridges (Waters Corp.). In brief, peptides were loaded onto HLB cartridge, washed with 0.1% trifluoroacetic acid (TFA), and eluted with 30% ACN and 60% ACN in 0.1% TFA. Flow through from the Oasis HLB was then loaded onto a Sep Pak C18 cartridge (Waters), washed with 0.1% TFA, and then eluted with 30% ACN and 60% ACN in 0.1% TFA. Eluates from HLB and C18 cartridges were combined and lyophilized in a vacuum centrifuge for LC-MS/MS proteome analysis as described below. *E. coli* cells were incubated at 37 °C with 200 rpm shaking and harvested at mid-log phase. Cells were pelleted at 10,000 x g, and then washed three times with phosphate-buffered saline pH 7.5. Harvested cell pellets were digested using the methods mentioned above.

### High-pH Reversed-phase Liquid Chromatography Fractionation

A total of 200 µg of in-house-prepared human and *E. coli* digests and Promega yeast protein digest were each fractionated by offline reversed-phase LC (Waters Acquity UPLC Peptide BEH300 C_18_ 1.7 µm, 1 mm i.d. x 150 mm column employing an Ultimate 3000 HPLC, Thermo Fisher Scientific) operating with column compartment temperature of 60 °C and flow rate of 100 µL/min. Mobile phase A consisted of 10 mM ammonium hydroxide, and mobile phase B consisted of 10 mM ammonium hydroxide/90% ACN. Sample loading and peptide separation were performed by applying a mixture of mobile phases as follows: i) 1% mobile phase B for 4 min, ii) 1% to 6% mobile phase B in 6 min, iii) 6% to 30% mobile phase B in 22 min, iv) 30% to 60% mobile phase B in 5 min, and v) ramp to 95% mobile phase B in 3 min. The washing step at 95% mobile phase B lasted for 2 min and was followed by an equilibration step at 1% mobile phase B for 5 min. Fractions were collected between 4 and 40 min; 36 fractions were collected for the human digest and 12 fractions were collected for yeast and *E.coli* protein digests. Each fraction was evaporated to dryness and resuspended by adding 10 µL or 30 µL for human and for yeast or *E. coli*, respectively, of an aqueous solution containing 0.1% (v/v) formic acid supplemented with the iRT kit (Biognosys) at a ratio 1:10 v/v.

### Analyses of Protein Digest Fractions by LC-MS/MS in DDA Mode to Support Spectral Library Generation

With the standardized and harmonized HRMS1 DIA workflows, the entire HRMS1-DIA data sets were acquired on the Q Exactive HF MS instrument at the eleven laboratories respectively. While all the spectral libraries were generated centrally via acquiring DDA files of the fractionated sample on the orbitrap-based mass spectrometry using HCD MS/MS fragmentation. The same capillary LC setup and separation conditions were applied for the HRMS1-DIA analysis, as well as the DDA analysis.

Each resuspended fraction of protein digest was analyzed with an Easy-nLC 1200 (Thermo Fisher Scientific) coupled to Orbitrap Fusion Lumos or Q Exactive HF-X mass spectrometers (Thermo Fisher Scientific) operated with DDA methods; 2 µL of each fraction was injected. Peptide separations were carried out on an Acclaim PepMap RSLC C_18_, 2 µm, 100 Å, 150 µm i.d. x 150 mm, nanoViper EASY-Spray column (Thermo Fisher Scientific). The column temperature was maintained at 50 °C using the EASY-Spray oven. Mobile phase A consisted of HPLC-grade water with 0.1% (v/v) formic acid, and mobile phase B consisted of HPLC-grade ACN with 20% (v/v) HPLC-grade water and 0.1% (v/v) formic acid. Samples were loaded at 3 µL/min with 100% mobile phase A for 2 min. Peptide elution was performed at 1.2 µL/min using the following gradient: i) 3% to 8% mobile phase B in 4 min, ii) 8% to 25% mobile phase B in 50 min, and iii) ramp to 80% mobile phase B in 4 min. The washing step at 80% mobile phase B lasted 2 min and was followed by an equilibration step at 100% A (1.7 min at 3 µL/min).

The Orbitrap Fusion Lumos mass spectrometer was configured for DDA using the full MS-data-dependent MS/MS setup and was operated in positive polarity mode. Spray voltage was set at 2 kV, funnel RF level at 40, and capillary temperature at 250 °C. Full MS survey scans were acquired at a resolution of 60,000 with an automatic gain control (AGC) target value of 4e5 and a maximum injection time of 20 ms over a scan range of *m/z* 350-1500. A data-dependent top 40 method was used during which up to 40 precursor ions were selected from each full MS scan to be fragmented through higher energy collisional dissociation (HCD). HCD MS/MS scans were acquired with a normalized collision energy of 30 at a resolution of 15,000 and with a starting mass of *m/z* 130. Precursor ions were isolated in a 1.6-Th window and accumulated to reach an AGC target value of 5e4 with a maximum injection time of 30 ms. Precursor ions with a charge state between 2 and 7 were selected for fragmentation, and the monoisotopic peak was isolated. Precursor ions and their isotopes selected for fragmentation were dynamically excluded for 30 s.

The Q Exactive HF-X mass spectrometer was configured for DDA using the full MS-data-dependent MS/MS setup and was operated in positive polarity mode. Spray voltage was set at 2 kV, funnel RF level at 40, and capillary temperature at 250 °C. Full MS survey scans were acquired at a resolution of 60,000 with an AGC target value of 3e6, a maximum injection time -of 20 ms, and a scan range of *m/z* 350-1500. A data-dependent top 20 method was used during which up to 20 precursor ions were selected from each full MS scan to be fragmented through HCD. HCD MS/MS scans were acquired with normalized collision energy 27 at a resolution of 15,000 with a starting mass of *m/z* 120. Precursor ions were isolated in a 1.6-Th window and accumulated to reach an AGC target value of 5e4 with a maximum injection time of 45 ms. Precursor ions with a charge state higher than 1 were selected for fragmentation, and the monoisotopic peak was isolated. Precursor ions and their isotopes selected for fragmentation were dynamically excluded for 40 s.

### Analyses of Mixed Proteomes Sampled by LC-MS/MS in DIA Mode

QC standards and Samples A and B were analyzed with an Easy-nLC 1200 or an Ultimate 3000 RSLCnano equipped with capillary flow meter coupled to a Q Exactive HF mass spectrometer (ThermoFisher Scientific) operated with DIA methods. Method settings are provided in Supplementary Protocols 2 and 3.

Peptide separations were carried out on an Acclaim PepMap RSLC C_18_, 2 µm, 100 Å, 150 µm i.d. x 150 mm, nanoViper EASY-Spray column (Thermo Fisher Scientific). The column temperature was maintained at 50 °C using the EASY-Spray oven. Mobile phase A consisted of HPLC-grade water with 0.1% (v/v) formic acid, and mobile phase B consisted of HPLC-grade ACN with 20% (v/v) HPLC-grade water and 0.1% (v/v) formic acid.

For LC-MS/MS analyses performed on the Easy-nLC 1200, samples were loaded at 4 µL/min with 100% mobile phase A for 5 min. Peptide elution was performed using the following gradient: i) 2% to 8% mobile phase B in 4 min, ii) 8% to 32% mobile phase B in 49 min, iii) 32% to 60 % mobile phase B in 1 min, and iv) ramp to 98% mobile phase B in 1 min at 2 µL/min. The washing step at 98% mobile phase B lasted 10 min (at 2 µL/min) and was followed by an equilibration step at 100% mobile phase A (6.7 min at 3 µL/min). After sample injection, the autosampler was washed by three cycles of “drawing/dispensing” 22 µL of HPLC-grade ACN containing 20% (v/v) HPLC-grade water and 0.1% (v/v) formic acid, followed by three cycles of “drawing/dispensing” 22 µL of HPLC-grade water containing 0.1% (v/v) formic acid.

For LC-MS/MS analyses performed on the Ultimate 3000 RSLCnano, samples were loaded at 3 µL/min with 100% mobile phase A for 5 min. Peptide elution was using the following gradient: i) 2% to 8% mobile phase B in 4 min, ii) 8% to 32% mobile phase B in 49 min, iii) 32% to 60% mobile phase B in 1 min, and iv) ramp to 98% mobile phase B in 1 min at 3 µL/min. After 5 min of run time, the inject valve was switched to the “load” position, and the autosampler procedure was triggered at 8.1 min. This allowed the injection loop to be washed and filled with a 20 µL plug of HPLC-grade ACN containing 20% (v/v) HPLC-grade water and 0.1% (v/v) formic acid. At 60 min, the injection valve was switched back to the “inject” position, allowing the 20 µL plug of ACN contained in the injection loop to be delivered quickly to the column for thorough washing. Concurrently (*i.e.*, at 60 min) the pump settings were modified to deliver 100% mobile phase A in the flow path for 13 min at 3 µL/min, which is sufficient time to achieve equilibration.

The Q Exactive HF mass spectrometer was configured for DIA by combining two experiment elements, corresponding to a full MS experiment and an MS/MS experiment, and was operated in positive polarity mode. Spray voltage was set in the range of 2 -2.4 kV to sustain a stable spray, funnel RF level was set at 50, and capillary temperature was maintained at 250 °C. The full MS experiment included one broadband scan acquired over *m/z* 400-1210 at a resolution of 120,000 with an AGC target value of 3e6 and with maximum injection time of 50 ms. The MS/MS experiment included 18 scans/cycle (for a total of 54 scans) acquired at R=30,000 with an AGC target value of 1e6 and with “Auto” maximum injection time. The precursor ions were isolated within a 15-Th window and fragmented by HCD acquired with normalized collision energy 28 and default charge state 3 and with a starting mass of *m/z* 200. Center values of isolation windows are reported in **Supplementary Protocols 2 and 3**.

### Search of DDA Data and Spectral Library Generation

The assignment of MS/MS spectra generated from DDA analyses of protein digests was made with Proteome Discoverer 2.2 software and Sequest HT algorithm using UniProt database filtered for “*Homo sapiens*” (downloaded April 2016), “*Saccharomyces cerevisiae*” (downloaded May 2016), or “*Escherichia coli*” (downloaded February 2016) taxonomies, concatenated with iRT peptide .fasta file (downloaded from the Biognosys webpage). Tolerances on precursors and fragment ions were set at +/- 10 ppm and +/- 0.02 Da, respectively. The searches were performed by specifying “Trypsin (full)” enzyme digest specificity constraints with a maximum of two missed cleavage sites allowed, “Oxidation” as dynamic modification, and “Carbamidomethylation” as static modification. The data were also searched against a decoy database, and the results were used to estimate q values using the Percolator algorithm within the Proteome Discoverer version 2.2 suite. Proteome discoverer result files were imported into Spectronaut Pulsar 11.0.15038.23.24843 (Asimov) software for the generation of the spectral libraries (.kit files) for each organism using default settings.

### Searching of DIA Data from Controlled Samples and Protein-Level Roll-up

A step-by-step procedure for data processing and evaluation with Spectronaut software is reported in **Supplementary Protocol 4**. Briefly, raw files (including QC standard and Sample A and Sample B raw data) were imported into Spectronaut software without conversion and searched against pertinent spectral libraries. The extraction of data used dynamic MS1 and MS2 mass tolerances, dynamic window for extracted ion current extraction window, and a non-linear iRT calibration strategy. The identification was carried out using a kernel density estimator and FDR cut-off of 0.01 at precursor and protein levels. The extracted quantitative data relied on MS1 data and benefited from interference correction and a local cross-run normalization strategy. The data processing results were exported using two customized reports for peptide and protein identification and for further quantification processing using R scripts. The peptide customized report included EG.PrecursorId, PG.ProteinAccessions, PG.ProteinDescriptions, EG.Qvalue, PEP.Quantity, and PG.Quantity fields. The protein customized report included PG.FastaFiles, PG.ProteinGroups, and PG.Quantity fields. The scripts were applied to subsets of analyses, which were grouped together according to day and lab. Through these scripts, additional filtering steps were applied prior to the comparison of quantifications of proteins between samples A and B including the removal of peptides shared by different organisms and the removal of sub-optimal precursor ions of peptides detected under different charge states (*i.e.*, the precursor ions not showing the highest number of retained quantitative data across the series of analyses considered). For each retained peptide, the maximum number of inter-sample combinations between the replicated analyses with quantitative data available was determined. This estimation was expanded at the protein level by summing these numbers obtained for all surrogate peptides. The proteins were considered as reliably quantifiable in the experiment and retained for further processing when at least eight combinations were available. The actual relative quantification of retained proteins was performed by calculating the geometric median of the peptide pairwise ratios obtained from all inter-sample combinations.

### HRMS1-DIA Analyses of Ovarian Cancer Samples

FFPE OCCC and HGSOC tissues were obtained under an IRB-approved protocol from INOVA Fairfax Hospital (Falls Church, VA, USA). Tissue specimens from two patients for each disease histotype were sectioned (8 µm) by microtome onto polyethylene naphthalate membrane slides (Leica Microsystems) and hematoxylin-eosin stained. Laser microdissection was used to harvest sections into LC/MS grade water. Tissue harvests were transferred to a Pressure Cycling Technology microtube (Pressure Biosciences, Inc.) containing 20 mL of 100 mM triethylammonium bicarbonate (TEAB) and 10% ACN. The samples were incubated at 99 °C for 30 min. Tissues were digested by adding SMART trypsin (ThermoFisher Scientific Inc.) at a ratio of 1 mg per 30,000,000 µm^2^ tissue in a 2320EXT barocycler (Pressure Biosciences, Inc.) where each sample was cycled 60 times between 45,000 psi for 50 s followed by 10 s at atmospheric pressure at a constant temperature of 50 °C. Each protein digest was transferred to a clean 0.5-mL microcentrifuge tube and lyophilized to ∼60% of total volume. Peptides digests were desalted (Thermo Scientific Pierce C_18_ Spin Columns), and peptide concentration was determined using the BCA assay. Equivalent amounts of tissue digests were shipped to three analytical laboratories, (*i.e.*, Lab 2, Lab 5, and Lab 9). Samples were resuspended to final concentrations of 1 µg/µL in 0.1% formic acid with iRT and analyzed in triplicate on a Q Exactive HF using the HRMS1-DIA method as described above. QC standards were analyzed in triplicate before and after cancer tissue sample analyses and a single QC standard analysis was performed midway through the overall analysis. For tissue-specific spectral library generation, 120 µg of total peptide digests were combined and fractionated by high pH reverse-phase liquid chromatography into 96 fractions using a linear gradient of ACN (0.69% per minute) as described above. Concatenated fractions were pooled into 36 fractions, lyophilized, and resuspended in 0.1% formic acid for analysis. Fractions were analyzed on a Q Exactive HF-X, and data were searched for spectral library generation as described above for HeLa cells. Tissue-specific and HeLa cell spectral libraries were combined in Spectronaut for analyses of HRMS1-DIA data. Protein abundance was determined for matched, histotype-specific HRMS1-DIA data collected by each analytical site. The proteins retained to undergo quantification were selected using the strategy described above but with the additional need to satisfy the criteria for the two histotype groups and at each analytical site. The abundance of retained proteins was estimated from each analysis by summing the intensities of individual peptides and averaging then across triplicates at each analytical site. Log_2_ fold-change protein abundances reflective of summed protein abundance ratios calculated relative to average protein abundances were quantified across all tissue samples for a given protein group. Differential analysis was performed using the LIMMA package (version 3.8) in R (version 3.5.2), and cluster analyses was performed using ClustVis (https://biit.cs.ut.ee/clustvis/). Spearman rho was calculated with features co-altered relative to a 112 protein and gene signature for frozen tissue stratifying high-grade serous and clear-cell ovarian cancers^28^, and box plots of relative standard deviation calculated from summed protein abundances corresponding to eighteen HGSOC/OCCC signature proteins of interest were performed using MedCalc (version 19.0.3).

## Data availability

The mass spectrometry proteomics data (.raw files) and spectral libraries used for the data processing (.kit files) have been deposited to the ProteomeXchange Consortium via the MassIVE partner repository with the dataset identifier MSV000084976

## Contributions

Y.X. directed the overall study design.

T.P.C. and N.W.B. designed the ovarian cancer study. Y.X., S.Ga., and Y.Z. contributed to protocol preparation.

Y.Z., B.H. and K.A. contributed to sample preparation and spectral libraries generation.

S.Go., B.H., C.L., R.B.K., S.P., D.C., H.Z., G.H., M.T., A.M., A.K., X.S., B.C., A.S., Y.J.C., C.R.J., H.Z.,

S.L., M.R.L., T.K., Z.C., B.L.P., and S.J.C. acquired the data, did data analysis onsite, and commented on the manuscript.

Y.X., N.W.B., S.Ga., P.N., N.P. and M.H. analyzed the data. Y.X., N.W.B., S.Ga., S.Go. and B.L.P prepared the figures

Y.X., N.W.B., S.Ga., S.Go., B.W. and T.P.C. wrote the manuscript.

## Acknowledgements

We would like to thank to Jenny Ho and Joshua J. Nicklay for evaluating and discussion about the study design, and for acquiring the data sets to establish the quality control criteria for the study; Oleksandr Boychenko for discussion about the capillary LC setup; We thank Steve Bios, Ming-yi He, and Scott M. Peterman for instrument maintenance and support with the MS measurements. David A. Joyce to setup ThemoCloud accounts to facilitate the data exchanges across the study; Sven Klingel and Shen Luan for assistance with the project coordination; Lukas Reiter for discussion about normalization and data analysis in the Spectronaut; Christoph Henrich and Alexander Tiegel for discussion about the data visualization; David Sarracino for the discussion about the study design. The views expressed herein are those of the authors and do not reflect the official policy of the Department of Army/Navy/Air Force, Department of Defense, Henry M. Jackson Foundation for the Advancement of Military Medicine, or U.S. Government.

## Supplementary Info

Supplementary Table 1: QC-DIA benchmark proteins quantified by analytical site.

Supplementary Table 2: MultiSite proteins quantification results of controlled samples (Log2 ratio protein abundance (Sample A / Sample B).

Supplementary Table 3: Tumor tissue proteins quantified by analytical site.

Supplementary Table 4: Significantly altered/ histotype correlated protein alterations.

Supplementary Table 5: The relative standard deviation (RSD) of 18 HGSOC and OCCC signature proteins across three analytical sites for each patient tissue sample.

Supplementary Protocol 1: Standard operating procedure - Sample preparation

Supplementary Protocol 2: Standard operating procedure - DIA Analysis with Capillary-flow UltiMate 3000 RSLCnano

Supplementary Protocol 3: Standard operating procedure - DIA Analysis with Easy nLC 1200

Supplementary Protocol 4: Standard operating procedure - DIA Analyses and Data Evaluation

