## Supplementary material for "Standardization and Harmonization of Distributed Multi-National Proteotype Analysis supporting Precision Medicine Studies": Standard operating procedure - Sample preparation

### 1) Material

| Description | Source | Product Number |
|---|---|---|
| Pierce HeLa Protein Digest Standard | Thermo Scientific™ | 88329 |
| MassPREP E. Coli Digest Standard | Waters | 186003196 |
| Mass Spec-Compatible Yeast Digest | Promega | V7461 |
| iRT-standard | Biognosys | Ki-3002-2 |
| 0.1% FA in Water, OPTIMA LC/MS | Fisher Chemicals | LS118-500 |
| *Protein LoBind Tubes, 0.5 mL, PCR clean, 100 tubes | Eppendorf | 30108094 |
| *Microvials PP, 0.3ml with short thread | VWR | 548-0440 |
| *Screw cap PP blue 9mm | VWR | 548-0088 |

*Recommended tubes and LC vials (and caps). Other low binding tubes and vials can be used as well.

### 2) Mixtures preparation

*Stock solutions*

iRT mixture (stock solution) (based on supplier instructions)
- Add 50 µL dissolution buffer (blue cap) to the iRT standard tube (red cap)
- Vortex the iRT standard tube for at least 1 minute.
- Subject tube to 5 min ultrasonic bath
- Store iRT mixture (stock solution) at 2-8°C (stable for 12 weeks)

HeLa digest stock solution (1 µg/µ - 100 µL)
- Defrost 10 vials of 20 µg of Hela digest (88329, lyophilized) at room temperature for 15 min
- Add 20 µL $H_20$ (+0.1% HCOOH) to each vial
- Vortex each vial for at least 1 minute
- Subject each vial to 5 min ultrasonic bath
- Vortex each vial for at least 30 s
- Keep one vial to prepare stock solution and transfer the content of 9 other vials in it (200 µL in total)

**Cancer Moonshot Multiple Sites Study**

Yeast digest stock solution (1 µg/µ - 100 µL)

- Defrost 1 vial of 100 µg Yeast digest (V7461, lyophilized) at room temperature for 15 min
- Add 100 µL $H_20$ (+0.1% HCOOH) to the vial
- Vortex the vial for at least 1 minute
- Subject the vial to 5 min ultrasonic bath
- Vortex the vial for at least 30 s
- Aliquot the stock solution (concentration 1µg/µL) into 50 µL volumes (0.5 mL low binding tubes) and freeze for future use.

E.Coli digest stock solution (1 µg/µ - 100 µL)

- Defrost 1 vial of 100 µg E.Coli digest (186003196, lyophilized) at room temperature for 15 min
- Add 100 µL $H_20$ (+0.1% HCOOH) to the vial
- Vortex the vial for at least 1 minute
- Subject the vial to 5 min ultrasonic bath
- Vortex the vial for at least 30 s
- Aliquot the stock solution (concentration 1µg/µL) into 25 µL volumes (0.5 mL low binding tubes) and freeze for future use.

*Multiple sites samples*

Mix A (1 µg/µL - 80 µL)

- Transfer 52 µL from Hela digest stock solution at 1µg/µL to a LC vial
- Transfer 12 µL from Yeast digest stock solution at 1µg/µL to the LC vial
- Transfer 16 µL from E. Coli digest stock solution at 1µg/µL to the LC vial
- Transfer 2.5 µL from iRT mixture (stock solution) to the LC vial
- Vortex to mix
- Put LC vial in autosampler. To be injected: 2µL/analysis (at least 30 µL needed for the study)

Mix B (1 µg/µL - 80 µL)

- Transfer 52 µL from Hela digest stock solution at 1µg/µL to a LC vial
- Transfer 24 µL from Yeast digest stock solution at 1µg/µL to the LC vial
- Transfer 4 µL from E. Coli digest stock solution at 1µg/µL to the LC vial
- Transfer 2.5 µL from iRT mixture (stock solution) to the LC vial
- Vortex to mix
- Put LC vial in autosampler. To be injected: 2µL/analysis (at least 30 µL needed for the study)

QC sample (1 µg/µL – 80-96 µL)

- Transfer the remaining content (should be between 80 and 96 µL) from Hela digest stock solution at 1µg/µL to a LC vial
- Transfer 3 µL from iRT mixture (stock solution) to the LC vial
- Vortex to mix
- Put LC vial in autosampler. To be injected: 2µL/analysis (at least 44 µL needed for the study)

Blank

- Transfer 300 µL $H_20$ (+0.1% HCOOH) to a LC vial
- Put LC vial in autosampler. To be injected: 2µL/analysis (at least 160 µL needed for the study)

### 3) Samples analyses plan

| Replicates | Day 1 | Day 2 | Day 3 | Day 4 | Day 5 | Day 6 | Day 7 |
|---|---|---|---|---|---|---|---|
| Blank | 1 | 1 | 1 | 1 | 1 | 1 | 1 |
| QC sample | 3 | 1 | 3 | 1 | 3 | 1 | 3 |
| Blank | 1 | 1 | 1 | 1 | 1 | 1 | 1 |
| Sample A | 3 | 1 | 3 | 1 | 3 | 1 | 3 |
| Blank | 1 | 1 | 1 | 1 | 1 | 1 | 1 |
| Sample B | 3 | 1 | 3 | 1 | 3 | 1 | 3 |
| Blank | 1 | 1 | 1 | 1 | 1 | 1 | 1 |
| QC sample | 1 | 1 | 1 | 1 | 1 | 1 | 1 |
| Blank (for 24 h total instrument time) | 4 (up to) | 10 (up to) | 4 (up to) | 10 (up to) | 4 (up to) | 10 (up to) | 4 (up to) |

### 4) Sample preparation summary

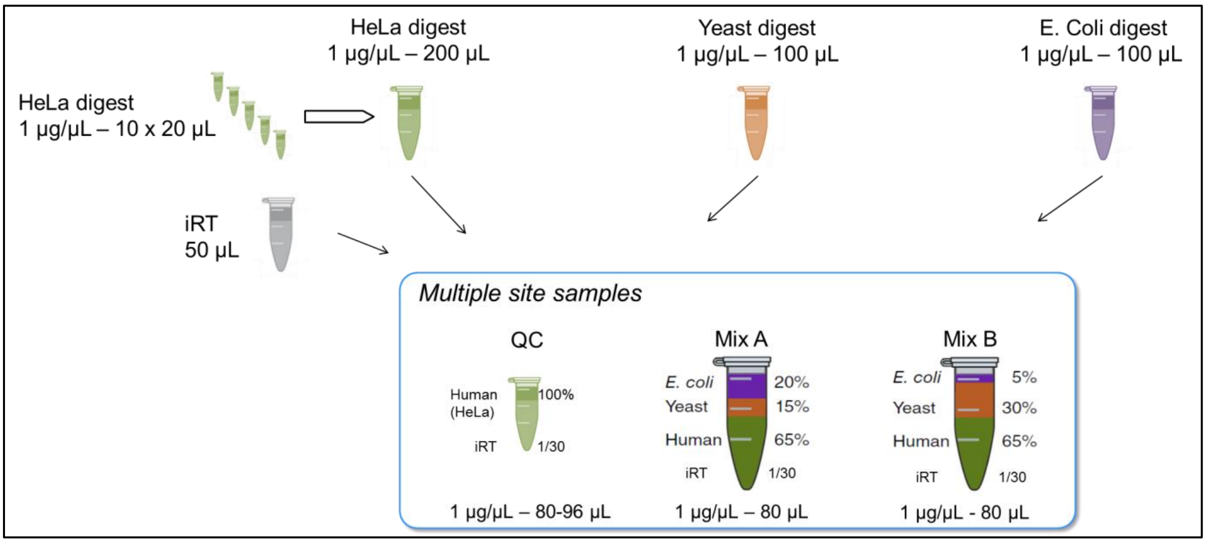
