## Supplementary material for "Standardization and Harmonization of Distributed Multi-National Proteotype Analysis supporting Precision Medicine Studies": Standard operating procedure - DIA Analysis with Capillary-flow UltiMate 3000 RSLCnano

### DIA Analysis – Capillary-flow UltiMate 3000 RSLCnano ES806

#### 1) Hardware configuration

- Mass Spectrometer: Q Exactive HF.

- Chromatographic System: UltiMate 3000 RSLCnano with capillary flow meter operated in a one-column setup.

Layout:

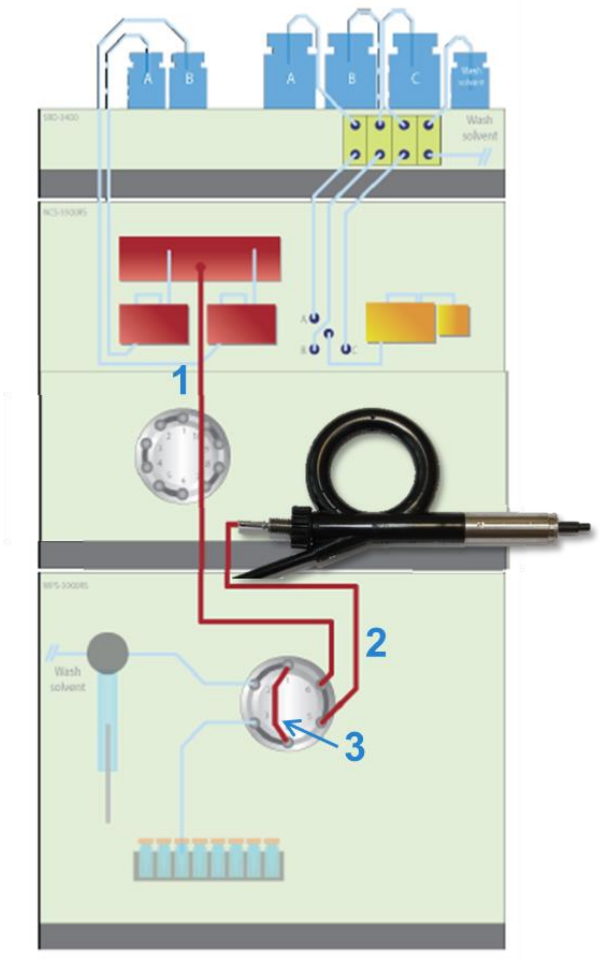

| # | Part | PN |
|---|---|---|
| 1 | nanoViper capillary FS/PEEK sheathed 1/32'' I.D. x L 50 µm x 550 mm | 6041.5560 |
| 2 | nanoViper capillary FS/PEEK sheathed 1/32'' I.D. x L 50 µm x 750 mm | 6041.5580 |
| 3 | nanoViper sample loop 20 µL, FS/PEEK sheathed | 6826.2420 |
| | EasySpray column PepMap RSLC $C_{18}$ 2 µm, 100A, 150 µm x 15 cm | ES806 |

#### 2) Instrument control software configuration

- 32bit PC: Foundation 3.1, Xcalibur 3.1, SII 1.2, Exactive 2.8SP1.
- 64 bit PC: Foundation 3.1 SP3 or SPE4, Xcalibur 4 or 4.1, SII 1.3, Exactive 2.8SP1 or Exactive 2.9.

#### 3) Samples and solvents

| Description | Source | Product Number / Reference |
|---|---|---|
| "Blank sample" - 0.1% FA in Water | Protocol "Part-I_Sample_Preparation" | Blank |
| "QC sample"<br>Human 100%<br>iRT 1/30 | Protocol "Part-I_Sample_Preparation" | QC – 1µg/µL |
| Mixed Proteomes "Sample A"<br>E. Coli 20%<br>Yeast 15%<br>Human 65%<br>iRT 1/30 | Protocol "Part-I_Sample_Preparation" | Mix A – 1 µg/µL |
| Mixed Proteomes "Sample B"<br>E. Coli 5%<br>Yeast 30%<br>Human 65%<br>iRT 1/30 | Protocol "Part-I_Sample_Preparation" | Mix B – 1 µg/µL |
| 0.1% FA in Water, OPTIMA LC/MS<br>- LC pump - solvent A | Fisher Chemicals | LS118-500 |
| 0.1% FA in 80 % Acetonitrile, OPTIMA LC/MS<br>- LC pump - solvent B<br>- Wash solvent | Fisher Chemicals | LS122-500 |
| Pierce™ LTQ Velos ESI Positive Ion Calibration Solution | Thermo Fisher Scientific | 88323 |
| *Microvials PP, 0.3ml with short thread | VWR | 548-0440 |
| *Screw cap PP blue 9mm | VWR | 548-0088 |

*Recommended tubes and LC vials (and caps). Other low binding tubes and vials can be used as well.

#### 4) LC and MS preparation and maintenance

Before launching the series of analyses, the LC-MS platform must be prepared through appropriate maintenance operations.

##### A) Liquid chromatography system

1. Prepare new solvents and subject them to ultrasonic bath for 15 min to remove dissolved gases.
2. Purge "Bothblocks" of NC pump for 30 min.
3. Purge "Flowmeter" of NC pump for 30 min.
4. Perform "Pressure Transducer Test" to verify that the offset of pressure transducers is correct and calibrate them if necessary.

5. Perform “Viscosity measurement” to verify the pump works correctly. The viscosity measured for channel A should be around 100% (+/- 5%). The viscosity measured for channel B should be around 60% (+/- 5%). Apply new viscosity values.
6. Perform syringe priming of autosampler using 10 cycles. Perform needle and fluidics washing with 50 µL of wash solvent.

**B) Mass spectrometer**

1. For testing if mass spectrometer is operating properly, infuse fresh calibration solution (product number 88323) into H-ESI source using syringe pump. Refer to manual “Q Exactive HF QuickStart Guide” for instructions about parameter settings to be used in Q Exactive HF Tune software for tuning and calibration (section “Getting Ions from Infusion Experiments”). Spray stability must be ≤ 10% (TIC Variation) to allow proper test and calibration (next steps).
2. Run “Isolation Transmission Endurance Test” (in “Extra Evaluation”). Cleaning of quadrupole is required for transmission score below 0.8 (while 1.0 is the optimal value).
3. Calibrate “Trapping Gas Control”. Delta pressure of the instrument should be below 5 bars.
4. Perform “Mass Calibration (pos)”.
5. Run all “Positive Ion Evaluation” procedure. Calibrate all parameters that did not pass the evaluation.

#### 5) LC method

- Solvent A: 0.1 % FA in Water.
- Solvent B: 0.1 % FA in 80 % Acetonitrile.
- Wash solvent: 0.1 % FA in 80 % Acetonitrile.
- Temperature (EASY- SPRAY source): 50°C.
- Injection mode: µLPickup. Transport vial(s) prepared by adding 300 µL of 0.1 % FA in water in Microvials (volume is sufficient for 18 analyses). Alternatively, transport vial(s) can be prepared by adding 5 mL of 0.1 % FA in water in 10 mL vials (PN 6820.0023).Transport vial must be changed (microvial) or clean/renewed (10 mL vials) every day or after a maximum of 18 injections. Position(s) in method have to match physical location.

- Gradient

| Time [min] | Flow [µL/min] | % B | Curve |
|---:|---:|---:|---:|
| - 13.000 | 3.000 | 2.0 | 5 |
| 5.000 | 3.000 | 2.0 | 5 |
| 9.000 | 1.200 | 8.0 | 5 |
| 58.000 | 1.200 | 32.0 | 5 |
| 59.000 | 3.000 | 60.0 | 5 |
| 60.000 | 3.000 | 98.0 | 5 |

- Commands added manually

| Time [min] | “Command” |
|---:|---:|
| - 13.000 | “Sampler.InjectValveToInject” |
| 6.000 | “Sampler. InjectValveToLoad” |
| 8.100 | “Sampler.Wash” |
| 60.000 | “Sampler.InjectValveToInject” |

- Complete LC method and screenshots are included in “9) Appendix – 1”

#### 6) MS parameters

- Tune Parameters

| Parameter | Value |
|---:|---:|
| Spray voltage [kV] | 2.00 (adjust +/-0.2 according to spray stability) |
| Capillary temperature [°C] | 250 |
| S-Lens RF level | 50 |

- MS Method

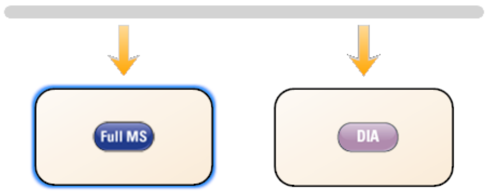

| **Parameter** | **Value** |
|---|---|
| ***Global settings*** | |
| use lock masses | off |
| Lock mass injection | - |
| Chrom. peak width (FWHM) | 15 s |
| ***Time*** | |
| Method duration | 60 min |
| ***Customized Tolerances (+/-)*** | |
| Inclusion | - |
| Lock Masses | - |
| Exclusion | - |
| Neutral lost | - |
| Mass Tag | - |

**Cancer Moonshot Multiple Sites Study**

| | |
|---|---|
| Dynamic Exclusion | - |
| ***Full MS*** | |
| Runtime | 0 to 60 min |
| Polarity | Positive |
| In-source CID | 0.0 ev |
| Microscans | 1 |
| Resolution | 120,000 |
| AGC target | 3e6 |
| Maximum IT | 50 ms |
| Number of scan ranges | 1 |
| Scan range | 400 – 1210 |
| Spectrum type | Profile |
| ***DIA*** | |
| Runtime | 0 to 60 min |
| Polarity | Positive |
| In-source CID | 0.0 ev |
| Default charge state | 3 |
| Microscan | 1 |
| Resolution | 30,000 |
| AGC target | 1e6 |
| Maximum IT | Auto |
| Loop count | 18 |
| MSX count | 1 |
| Isolation window | 15 m/z |
| Isolation offset | 0.0 m/z |
| Fixed first mass | 200 m/z |

| | |
|---|---|
| NCE/ stepped NCE | nce : 28 |
| Spectrum data type | Profile |

- Full inclusion list for DIA scans is included in "10) Appendix – 2"

#### 7) Samples analyses plan

- Samples analysis plan introduced in Supplementary Protocol 1 is reminded below.
- The .raw data files must be named under following rule "Lab_W-Day_X-Sample_Y-Rep_Z", where "W" is the lab number (*e.g.*, "1" or "2"), "X" is the day number (1-7), "Y" is the sample name ("Blank", "QC", "A", or "B"), and "Z" is the replicate number (1-3).
- Some examples of .raw data file names are included below:
    - "Lab_1-Day_1-Sample_Blank-Rep_2"
    - "Lab_1-Day_5-Sample_QC-Rep_1"
    - "Lab_2-Day_3-Sample_A-Rep_3"

| Replicates | Day 1 | Day 2 | Day 3 | Day 4 | Day 5 | Day 6 | Day 7 |
|---|---|---|---|---|---|---|---|
| Blank | 1 | 1 | 1 | 1 | 1 | 1 | 1 |
| QC sample | 3 | 1 | 3 | 1 | 3 | 1 | 3 |
| Blank | 1 | 1 | 1 | 1 | 1 | 1 | 1 |
| Sample A | 3 | 1 | 3 | 1 | 3 | 1 | 3 |
| Blank | 1 | 1 | 1 | 1 | 1 | 1 | 1 |
| Sample B | 3 | 1 | 3 | 1 | 3 | 1 | 3 |
| Blank | 1 | 1 | 1 | 1 | 1 | 1 | 1 |
| QC sample | 1 | 1 | 1 | 1 | 1 | 1 | 1 |
| Blank (for 24 h total instrument time) | 4 (up to) | 10 (up to) | 4 (up to) | 10 (up to) | 4 (up to) | 10 (up to) | 4 (up to) |

#### 8) General consideration for sample analyses and related data evaluation

- The data processing and evaluation procedure is described in details in Supplementary Protocol 4.
- In day 1, following initial blank analysis and before triplicated analyses of QC, it is recommended to carry-out a first analysis of QC sample in order to passivate brand new chromatography column. This initial analysis of QC sample must not be included in QC evaluation procedure.
- Each day, the evaluation of "blank" analysis must be performed before starting analyses of "QC sample". In case the evaluation criteria are not fulfilled, stop LC-MS/MS analyses sequence and

troubleshoot possible issues (refer to troubleshooting guides of instruments). Repeat failed analysis once the issue has been fixed.
- In days 1, 3, 5, and 7, the evaluation of “QC sample” analyses in triplicate must be performed before starting analyses of “Sample A”. In case the QC criteria are not fulfilled, stop LC-MS/MS analyses sequence and troubleshoot possible issues (refer to troubleshooting guide of instruments). Repeat failed analyses once the issue has been fixed.
- In general, the evaluation of data should be carried out in the course of the study to detect major instrument dysfunction as soon as possible and stop LC-MS/MS analyses sequence in order to avoid wasting samples.

#### 9) Appendix – 1: Complete LC method and screenshots

##### Complete LC method script :

- {Initial Time} Instrument Setup
- PumpModule.LoadingPump.%A.Equate "%A 0.1% FA"
- PumpModule.LoadingPump.%B.Equate "%B 0.1% FA ACN"
- PumpModule.LoadingPump.%C.Equate "%C"
- PumpModule.LoadingPump.Pressure.LowerLimit 0 [bar]
- PumpModule.LoadingPump.Pressure.UpperLimit 500 [bar]
- PumpModule.LoadingPump.MaximumFlowRampUp 998 [µl/min²]
- PumpModule.LoadingPump.MaximumFlowRampDown 998 [µl/min²]
- PumpModule.NC_Pump.%A.Equate "%A 0.1% FA"
- PumpModule.NC_Pump.%B.Equate "%B 0.1% FA in 80% ACN"
- PumpModule.NC_Pump.Pressure.LowerLimit 0 [bar]
- PumpModule.NC_Pump.Pressure.UpperLimit 900 [bar]
- PumpModule.NC_Pump.MaximumFlowRampUp 99.000 [µl/min²]
- PumpModule.NC_Pump.MaximumFlowRampDown 99.000 [µl/min²]
- ColumnOven.TempCtrl Off
- Sampler.LowDispersionMode Off
- Sampler.WashSpeed 4.000 [µl/s]
- Sampler.WashVolume 50.000 [µl]
- Sampler.PunctureDepth 6.000 [mm]
- Sampler.SampleHeight 1.000 [mm]
- Sampler.WasteSpeed 4.000 [µl/s]
- Sampler.DispenseDelay 2.000 [s]
- Sampler.DispSpeed 2.000 [µl/s]
- Sampler.DrawSpeed 0.200 [µl/s]
- Sampler.DrawDelay 5.000 [s]
- Sampler.RinseBetweenReinjections Yes
- Sampler.FlushVolume 6.000 [µl]
- Sampler.TransVialPunctureDepth 6.000 [mm]
- Sampler.TransLiquidHeight 3.000 [mm]
- Sampler.TransportVialCapacity 99999
- Sampler.LastTransportVial RA8
- Sampler.FirstTransportVial RA8
- Sampler.InjectMode ulPickUp
- Sampler.LoopWashFactor 2.000
- Sampler.PumpDevice "NC_Pump"
- Sampler.TempCtrl On
- Sampler.Temperature.Nominal 5.0 [°C]
- Sampler.ReadyTempDelta None
- Sampler.Temperature.LowerLimit 4.0 [°C]
- Sampler.Temperature.UpperLimit 45.0 [°C]
- -13.000 Equilibration Duration = 13.000 [min]
- Sampler.InjectValveToInject
- PumpModule.LoadingPump.Flow.Nominal 0.000 [µl/min]
- PumpModule.LoadingPump.%B.Value 0.0 [%]
- PumpModule.LoadingPump.%C.Value 0.0 [%]
- PumpModule.LoadingPump.Curve 5
- PumpModule.NC_Pump.Flow.Nominal 3.000 [µl/min]
- PumpModule.NC_Pump.%B.Value 2.0 [%]
- PumpModule.NC_Pump.Curve 5
- 0.000 Inject Preparation
- Wait PumpModule.LoadingPump.Ready And PumpModule.NC_Pump.Ready And ColumnOven.Ready And Sampler.Ready
- 0.000 Inject
- Sampler.Inject
- 0.000 Start Run
- ColumnOven.ColumnOven_Temp.AcqOn
- PumpModule.LoadingPump.LoadingPump_Pressure.AcqOn
- PumpModule.NC_Pump.NC_Pump_Pressure.AcqOn
- 0.000 Run Duration = 60.000 [min]
- PumpModule.LoadingPump.Flow.Nominal 0.000 [µl/min]
- PumpModule.LoadingPump.%B.Value 0.0 [%]
- PumpModule.LoadingPump.%C.Value 0.0 [%]
- PumpModule.LoadingPump.Curve 5
- 5.000
- PumpModule.NC_Pump.Flow.Nominal 3.000 [µl/min]
- PumpModule.NC_Pump.%B.Value 2.0 [%]
- PumpModule.NC_Pump.Curve 5
- 6.000
- Sampler.InjectValveToLoad
- 8.100
- Sampler.Wash
- 9.000
- PumpModule.NC_Pump.Flow.Nominal 1.200 [µl/min]
- PumpModule.NC_Pump.%B.Value 8.0 [%]
- PumpModule.NC_Pump.Curve 5

- 58.000
- PumpModule.NC_Pump.Flow.Nominal 1.200 [µl/min]
- PumpModule.NC_Pump.%B.Value 32.0 [%]
- PumpModule.NC_Pump.Curve 5
- 59.000
- PumpModule.NC_Pump.Flow.Nominal 3.000 [µl/min]
- PumpModule.NC_Pump.%B.Value 60.0 [%]
- PumpModule.NC_Pump.Curve 5
- 60.000
- Sampler.InjectValveToInject
- PumpModule.NC_Pump.Flow.Nominal 3.000 [µl/min]
- PumpModule.NC_Pump.%B.Value 98.0 [%]
- PumpModule.NC_Pump.Curve 5
- 60.000 Stop Run
- ColumnOven.ColumnOven_Temp.AcqOff
- PumpModule.LoadingPump.LoadingPump_Pressure.AcqOff
- PumpModule.NC_Pump.NC_Pump_Pressure.AcqOff
- End

### Complete LC method screenshots :

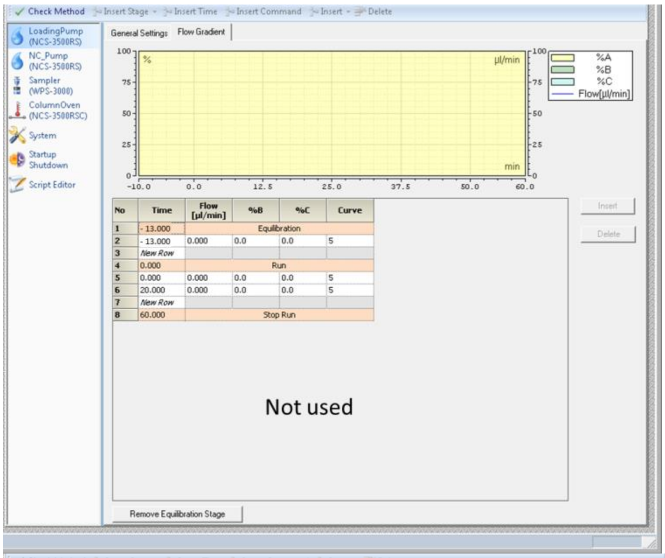

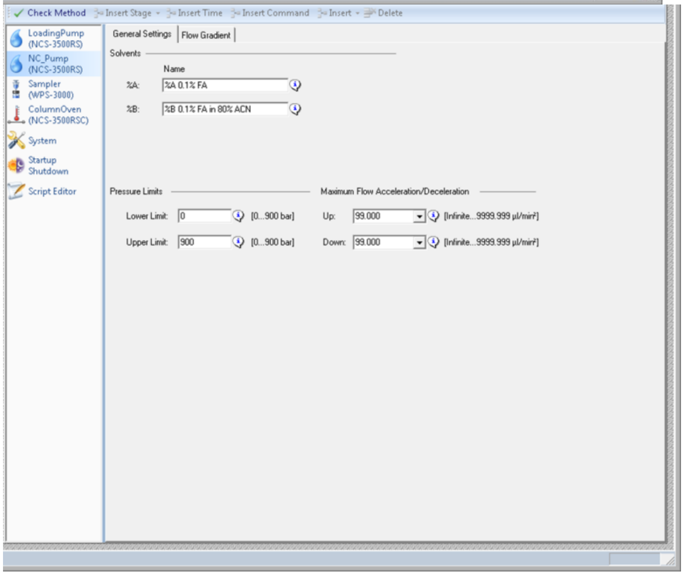

#### Cancer Moonshot Multiple Sites Study

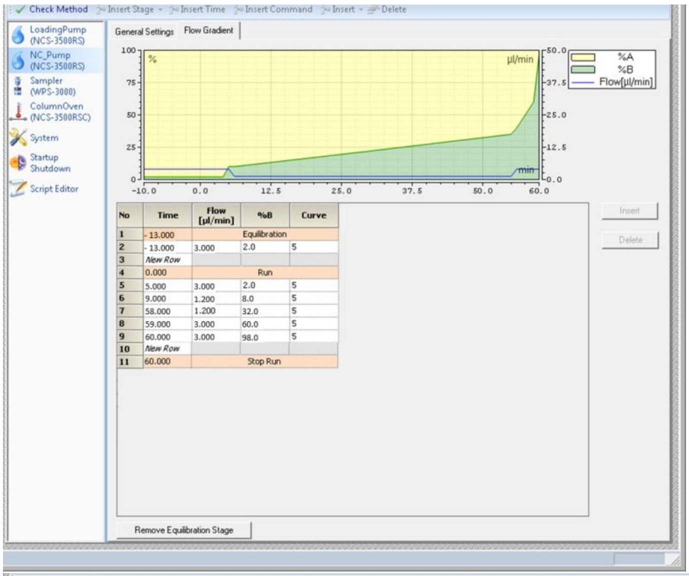

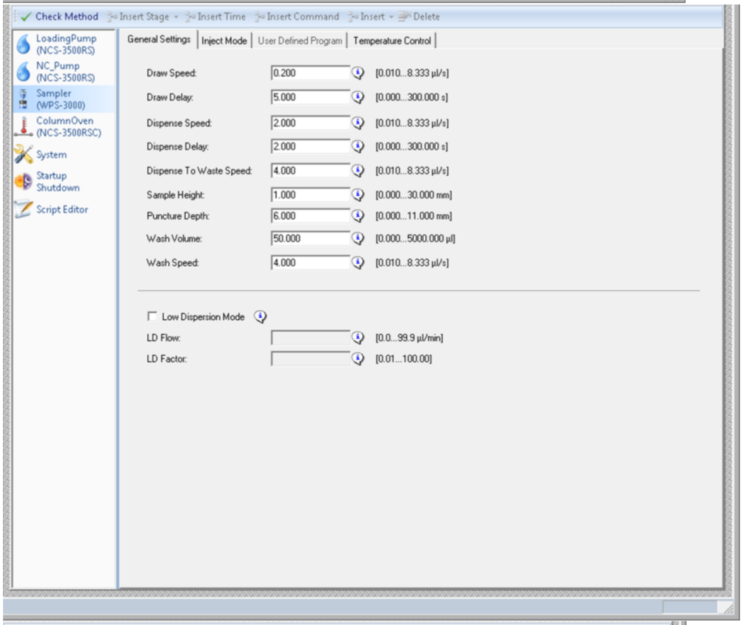

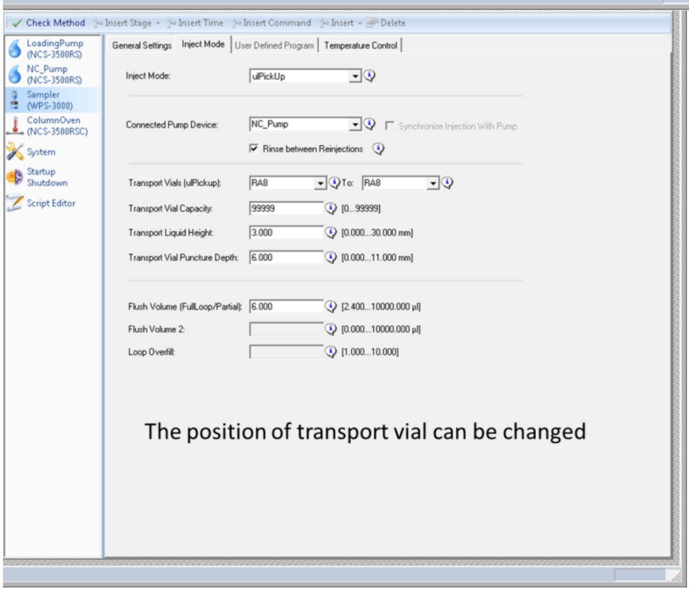

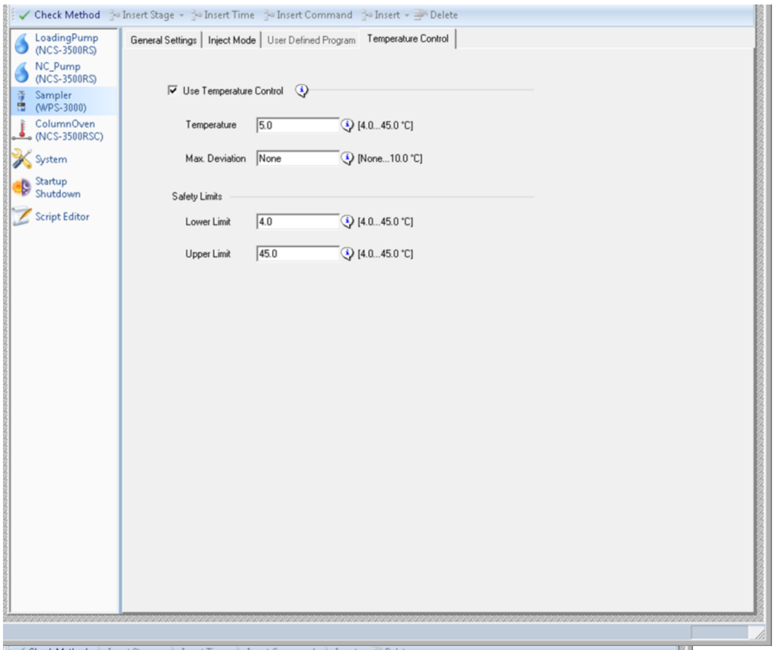
Check Method
Insert Stage
Insert Time
Insert Command
Insert
Delete
LoadingPump (NCS-3500RS)
NC_Pump (NCS-3500RS)
Sampler (WPS-3000)
ColumnOven (NCS-3500RSC)
System
Startup Shutdown
Script Editor
General Settings
Inject Mode
User Defined Program
Temperature Control
Use Temperature Control
Temperature
5.0
[4.0...45.0 °C]
Max. Deviation
None
[None...10.0 °C]
Safety Limits
Lower Limit
4.0
[4.0...45.0 °C]
Upper Limit
45.0
[4.0...45.0 °C]

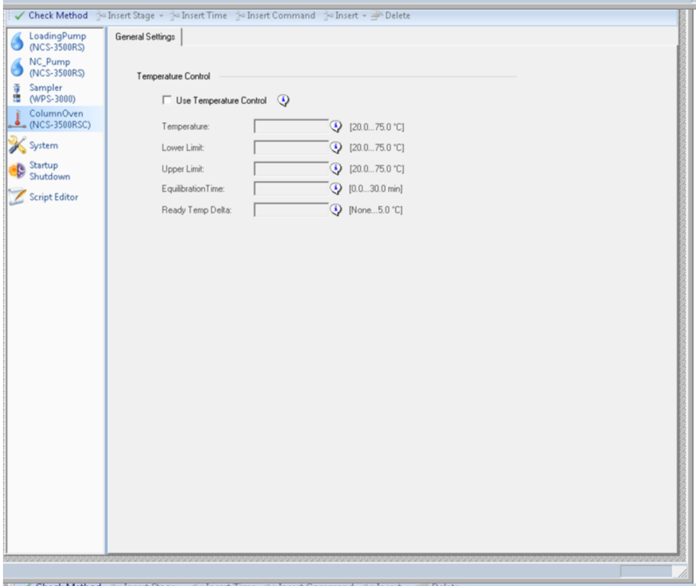
Check Method
Insert Stage
Insert Time
Insert Command
Insert
Delete
LoadingPump (NCS-3500RS)
NC_Pump (NCS-3500RS)
Sampler (WPS-3000)
ColumnOven (NCS-3500RSC)
System
Startup Shutdown
Script Editor
General Settings
Temperature Control
Use Temperature Control
Temperature:
[20.0...75.0 °C]
Lower Limit:
[20.0...75.0 °C]
Upper Limit:
[20.0...75.0 °C]
EquilibrationTime:
[0.0...30.0 min]
Ready Temp Delta:
[None...5.0 °C]

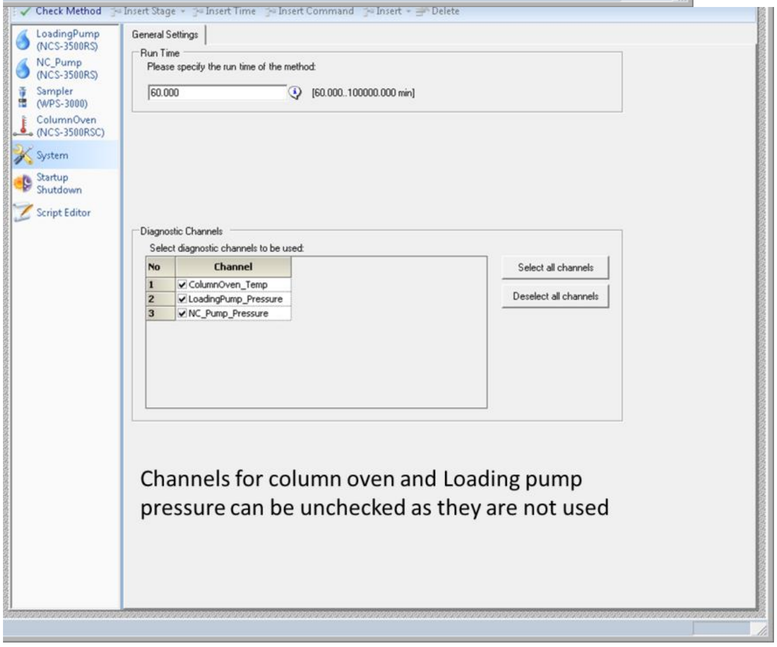
Check Method
Insert Stage
Insert Time
Insert Command
Insert
Delete
LoadingPump (NCS-3500RS)
NC_Pump (NCS-3500RS)
Sampler (WPS-3000)
ColumnOven (NCS-3500RSC)
System
Startup Shutdown
Script Editor
General Settings
Run Time
Please specify the run time of the method:
60.000
[60.000...100000.000 min]
Diagnostic Channels
Select diagnostic channels to be used:
No
Channel
1
ColumnOven_Temp
2
LoadingPump_Pressure
3
NC_Pump_Pressure
Select all channels
Deselect all channels
Channels for column oven and Loading pump pressure can be unchecked as they are not used

#### 10) Appendix – 2: Full inclusion list for DIA scans

| Mass [m/z] | Formula [M] | Formula type | Species | CS [z] | Polarity | Start [min] | End [min] | (N)CE | (N)CE type | MSX ID | Comment |
|---|---|---|---|---|---|---|---|---|---|---|---|
| 407.5 | | | | | Positive | | | | | | |
| 422.5 | | | | | Positive | | | | | | |
| 437.5 | | | | | Positive | | | | | | |
| 452.5 | | | | | Positive | | | | | | |
| 467.5 | | | | | Positive | | | | | | |
| 482.5 | | | | | Positive | | | | | | |
| 497.5 | | | | | Positive | | | | | | |
| 512.5 | | | | | Positive | | | | | | |
| 527.5 | | | | | Positive | | | | | | |
| 542.5 | | | | | Positive | | | | | | |
| 557.5 | | | | | Positive | | | | | | |
| 572.5 | | | | | Positive | | | | | | |
| 587.5 | | | | | Positive | | | | | | |
| 602.5 | | | | | Positive | | | | | | |
| 617.5 | | | | | Positive | | | | | | |
| 632.5 | | | | | Positive | | | | | | |
| 647.5 | | | | | Positive | | | | | | |
| 662.5 | | | | | Positive | | | | | | |
| 677.5 | | | | | Positive | | | | | | |
| 692.5 | | | | | Positive | | | | | | |
| 707.5 | | | | | Positive | | | | | | |
| 722.5 | | | | | Positive | | | | | | |
| 737.5 | | | | | Positive | | | | | | |
| 752.5 | | | | | Positive | | | | | | |
| 767.5 | | | | | Positive | | | | | | |
| 782.5 | | | | | Positive | | | | | | |

**Cancer Moonshot Multiple Sites Study**

| | | | | | |
|---|---|---|---|---|---|
| 797.5 | | | | | Positive |
| 812.5 | | | | | Positive |
| 827.5 | | | | | Positive |
| 842.5 | | | | | Positive |
| 857.5 | | | | | Positive |
| 872.5 | | | | | Positive |
| 887.5 | | | | | Positive |
| 902.5 | | | | | Positive |
| 917.5 | | | | | Positive |
| 932.5 | | | | | Positive |
| 947.5 | | | | | Positive |
| 962.5 | | | | | Positive |
| 977.5 | | | | | Positive |
| 992.5 | | | | | Positive |
| 1007.5 | | | | | Positive |
| 1022.5 | | | | | Positive |
| 1037.5 | | | | | Positive |
| 1052.5 | | | | | Positive |
| 1067.5 | | | | | Positive |
| 1082.5 | | | | | Positive |
| 1097.5 | | | | | Positive |
| 1112.5 | | | | | Positive |
| 1127.5 | | | | | Positive |
| 1142.5 | | | | | Positive |
| 1157.5 | | | | | Positive |
| 1172.5 | | | | | Positive |
| 1187.5 | | | | | Positive |
| 1202.5 | | | | | Positive |
