## Supplementary material for "Standardization and Harmonization of Distributed Multi-National Proteotype Analysis supporting Precision Medicine Studies": Standard operating procedure - DIA Analysis with Easy nLC 1200

### DIA Analysis – Easy nLC 1200 ES806

#### 1) Hardware configuration

- Mass Spectrometer: Q Exactive HF.

- Chromatographic System: Easy nLC 1200 operated in a one-column setup.

Layout:

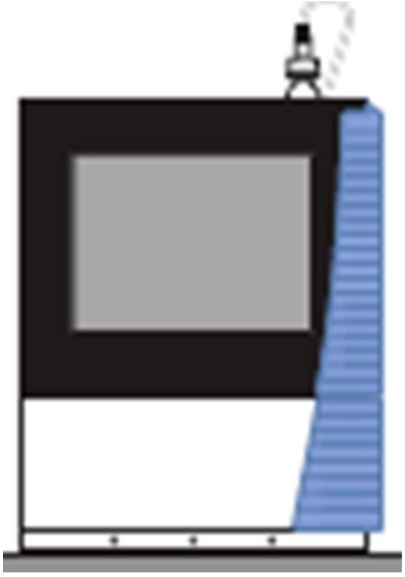

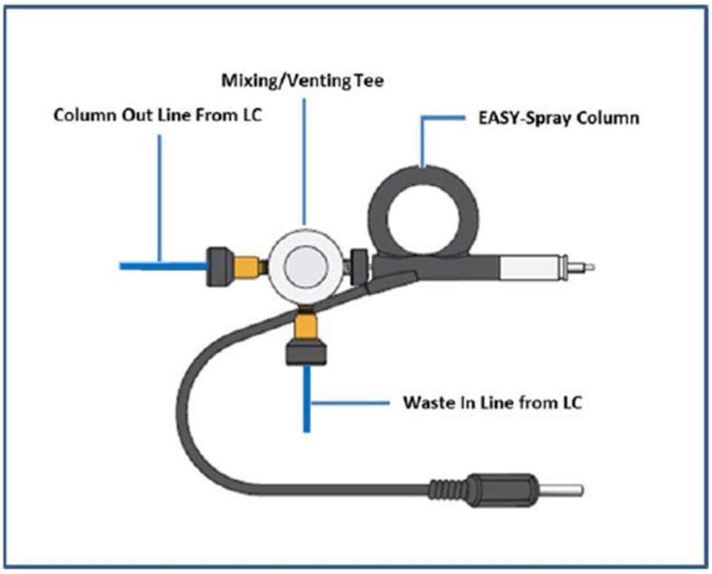

| Part | PN |
|---|---|
| nanoViper sample loop 20 µL, FS/PEEK sheathed | 6826.2420 |
| EASY-Spray column PepMap RSLC $C_{18}$ 2 µm, 100A, 150 µm x 15 cm | ES806 |
| Column out line from LC: nanoViper, 20 µm ID, 550 mm length | 6041.5261 (LC560) |

#### 2) Instrument control software configuration

- 32bit PC: Foundation 3.1, Xcalibur 3.1, Exactive 2.8SP1, EASY-nLC 1200 system: LC Devices 3.00.
- 64 bit PC: Foundation 3.1 SP3 or SPE4, Xcalibur 4 or 4.1, Exactive 2.8SP1 or Exactive 2.9, EASY-nLC 1200 system: LC Devices 3.00.

#### 3) Samples and solvents

| Description | Source | Product Number / Reference |
|---|---|---|
| "Blank sample" - 0.1% FA in Water | Protocol "Part-I_Sample_Preparation" | Blank |
| "QC sample"<br>Human 100%<br>iRT 1/30 | Protocol "Part-I_Sample_Preparation" | QC – 1µg/µL |
| Mixed Proteomes "Sample A"<br>E. Coli 20%<br>Yeast 15%<br>Human 65%<br>iRT 1/30 | Protocol "Part-I_Sample_Preparation" | Mix A – 1 µg/µL |
| Mixed Proteomes "Sample B"<br>E. Coli 5%<br>Yeast 30%<br>Human 65%<br>iRT 1/30 | Protocol "Part-I_Sample_Preparation" | Mix B – 1 µg/µL |
| 0.1% FA in Water, OPTIMA LC/MS<br>- LC pump - solvent A<br>- Wash solvent – bottle 3 | Fisher Chemicals | LS118-500 |
| 0.1% FA in 80 % Acetonitrile, OPTIMA LC/MS<br>- LC pump - solvent B<br>- Wash solvent – bottle 1 | Fisher Chemicals | LS122-500 |
| Pierce™ LTQ Velos ESI Positive Ion Calibration Solution | Thermo Fisher Scientific | 88323 |
| *Microvials PP, 0.3ml with short thread | VWR | 548-0440 |
| *Screw cap PP blue 9mm | VWR | 548-0088 |

*Recommended tubes and LC vials (and caps). Other low binding tubes and vials are fine as well.

#### 4) LC and MS preparation and maintenance

Before launching the series of analyses, the LC-MS platform must be prepared through appropriate maintenance operations.

##### A) Liquid chromatography system

1. Prepare new solvents and subject them to ultrasonic bath for 15 min to remove dissolved gases.
2. Purge solvent of pump A, pump B, and pump S using 5 iterations.

3. Flush air of pump A, pump B, and pump S until the required threshold volume of 10 µL is reached.
4. Calibrate flow sensors for A and B solvents if different solvent composition was used previously.

**B) Mass spectrometer**

1. For testing if mass spectrometer is operating properly, infuse fresh calibration solution (product number 88323) into H-ESI source using syringe pump. Refer to manual "Q Exactive HF QuickStart Guide" for instructions about parameter settings to be used in Q Exactive HF Tune software for tuning and calibration (section "Getting Ions from Infusion Experiments"). Spray stability must be ≤ 10% (TIC Variation) to allow proper test and calibration (next steps).
2. Run "Isolation Transmission Endurance Test" (in "Extra Evaluation"). Cleaning of quadrupole is required for transmission score below 0.8 (while 1.0 is the optimal value).
3. Calibrate "Trapping Gas Control". Delta pressure of the instrument should be below 5 bars.
4. Perform "Mass Calibration (pos)".
5. Run all "Positive Ion Evaluation" procedure. Calibrate all parameters that did not pass the evaluation.

#### 5) LC method

- Solvent A: 0.1 % FA in Water.
- Solvent B: 0.1 % FA in 80 % Acetonitrile.
- Wash solvent: 0.1 % FA in Water (bottle 3); 0.1 % FA in 80 % Acetonitrile (bottle 1).
- Temperature (EASY- Spray source): 50°C.
- Temperature autosampler: 5°C.

.

- Gradient

| Time [min] | Duration[min] | Flow rate [µL/min] | %B |
|---|---|---|---|
| 0.000 | N/A | 1 200 | 2.0 |
| 4.000 | 4.000 | 1 200 | 8.0 |
| 53.000 | 49.000 | 1 200 | 32.0 |
| 54.000 | 1.000 | 1 200 | 60.0 |
| 55.000 | 1.000 | 2 000 | 98.0 |
| 65.000 | 10.000 | 2 000 | 98.0 |

- Additional settings

| | Volume [µL] | Flow [µL/min] | Max. pressure [Bar] |
|---|---|---|---|
| Sample pickup | 2 | 10.00 | - |
| Sample loading | 20 | 4.00 | 1000.00 |
| Analytical column equilibration | 20 | 3.00 | 1000.00 |

- Screenshots of the complete method are included in "9) Appendix – 1"

#### 6) MS parameters

- Tune Parameters

| Parameter | Value |
|---|---|
| Spray voltage [kV] | 2.00 (adjust +/-0.2 according to spray stability) |
| Capillary temperature [°C] | 250 |
| S-Lens RF level | 50 |

- MS Method

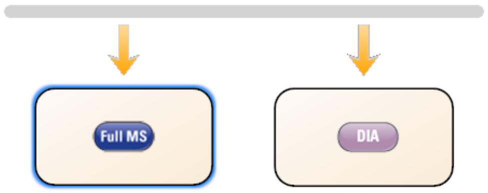

| Parameter | Value |
|---|---|
| ***Global settings*** | |
| use lock masses | off |
| Lock mass injection | - |
| Chrom. peak width (FWHM) | 15 s |
| ***Time*** | |
| Method duration | 60 min |
| ***Customized Tolerances (+/-)*** | |
| Inclusion | - |
| Lock Masses | - |
| Exclusion | - |
| Neutral lost | - |
| Mass Tag | - |
| Dynamic Exclusion | - |
| ***Full MS*** | |
| Runtime | 0 to 60 min |

#### 9) Appendix – 1: Complete LC method screenshots

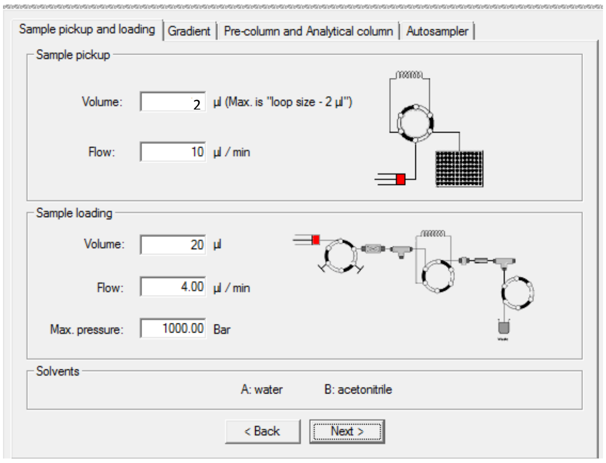

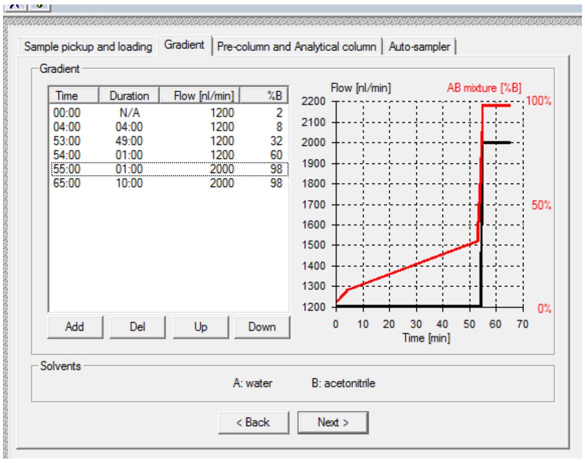

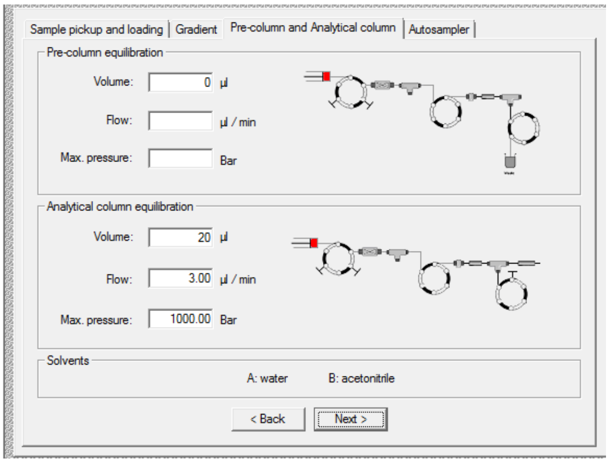

#### Cancer Moonshot Multiple Sites Study

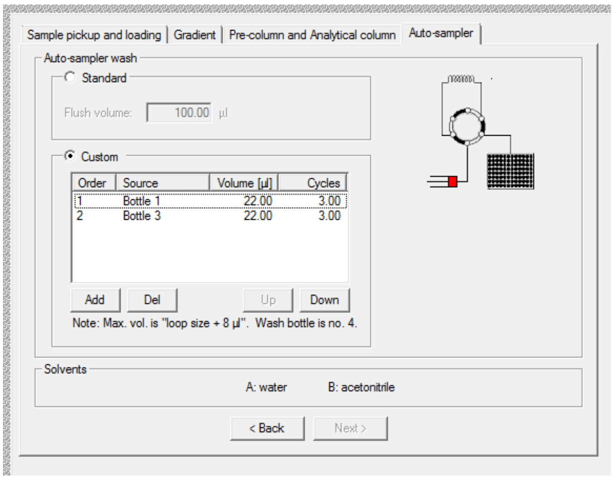

#### 10) Appendix – 2: Full inclusion list for DIA scans

| Mass [m/z] | Formula [M] | Formula type | Species | CS [z] | Polarity | Start [min] | End [min] | (N)CE | (N)CE type | MSX ID | Comment |
|---|---|---|---|---|---|---|---|---|---|---|---|
| 407.5 | | | | | Positive | | | | | | |
| 422.5 | | | | | Positive | | | | | | |
| 437.5 | | | | | Positive | | | | | | |
| 452.5 | | | | | Positive | | | | | | |
| 467.5 | | | | | Positive | | | | | | |
| 482.5 | | | | | Positive | | | | | | |
| 497.5 | | | | | Positive | | | | | | |
| 512.5 | | | | | Positive | | | | | | |
| 527.5 | | | | | Positive | | | | | | |
| 542.5 | | | | | Positive | | | | | | |
| 557.5 | | | | | Positive | | | | | | |
| 572.5 | | | | | Positive | | | | | | |
| 587.5 | | | | | Positive | | | | | | |
| 602.5 | | | | | Positive | | | | | | |
| 617.5 | | | | | Positive | | | | | | |
| 632.5 | | | | | Positive | | | | | | |
| 647.5 | | | | | Positive | | | | | | |
| 662.5 | | | | | Positive | | | | | | |
| 677.5 | | | | | Positive | | | | | | |
| 692.5 | | | | | Positive | | | | | | |
| 707.5 | | | | | Positive | | | | | | |
| 722.5 | | | | | Positive | | | | | | |
| 737.5 | | | | | Positive | | | | | | |
| 752.5 | | | | | Positive | | | | | | |
| 767.5 | | | | | Positive | | | | | | |
| 782.5 | | | | | Positive | | | | | | |

**Cancer Moonshot Multiple Sites Study**

| | | | | | |
|---|---|---|---|---|---|
| 797.5 | | | | | Positive |
| 812.5 | | | | | Positive |
| 827.5 | | | | | Positive |
| 842.5 | | | | | Positive |
| 857.5 | | | | | Positive |
| 872.5 | | | | | Positive |
| 887.5 | | | | | Positive |
| 902.5 | | | | | Positive |
| 917.5 | | | | | Positive |
| 932.5 | | | | | Positive |
| 947.5 | | | | | Positive |
| 962.5 | | | | | Positive |
| 977.5 | | | | | Positive |
| 992.5 | | | | | Positive |
| 1007.5 | | | | | Positive |
| 1022.5 | | | | | Positive |
| 1037.5 | | | | | Positive |
| 1052.5 | | | | | Positive |
| 1067.5 | | | | | Positive |
| 1082.5 | | | | | Positive |
| 1097.5 | | | | | Positive |
| 1112.5 | | | | | Positive |
| 1127.5 | | | | | Positive |
| 1142.5 | | | | | Positive |
| 1157.5 | | | | | Positive |
| 1172.5 | | | | | Positive |
| 1187.5 | | | | | Positive |
| 1202.5 | | | | | Positive |
