## Supplementary material for "Standardization and Harmonization of Distributed Multi-National Proteotype Analysis supporting Precision Medicine Studies": Standard operating procedure - DIA Analyses and Data Evaluation

### DIA Analyses Data Evaluation - Processing

#### I) General Considerations

- The samples analyses plan is provided in Section 3 of Supplementary Protocol 1.
- Sections V to VI in the document below describe the data processing / evaluation procedure for "Day 1" experiment. The procedure must be repeated for "Day 3", "Day 5, and "Day 7".
- Section VII describes the centralized data processing of controlled samples A and B analyses performed across the different lab and days retained.

#### II) Data processing software

- Biognosys Spectronaut Pulsar, version 11 and higher.
- Xcalibur, version 3.1 or higher.

#### III) Preliminary operations

1. Copy on processing computer spectral library files "ES806_KG_36F.kit" (Human), "yeast_ES806_12F.kit" (Yeast), and Ecoli_ES806_12F.kit (E. Coli).
2. Copy on processing computer protein databases "swissprot_homo_201604.fasta" (Human), "Saccharomyces cerevisiae_201602" (Yeast), and "Uniprot-Ecoli_K12_201709.fasta" (E. Coli).
3. Copy on processing computer report templates for proteins "quanBenchmark_protein.rs" and peptides "quanBenchmark_peptides.rs".

#### IV) Preparation in Spectronaut

1. Start the program "Spectronaut".
2. Select the tab: Prepare.
3. Press the link: Import Spectral Library.
4. In popup window, select Human spectral library file.
5. Press the button: Open.
6. Repeat steps IV – 3, 4, and 5 for Yeast spectral library, and then E. Coli spectral library.
7. Select the tab: Databases.
8. Select the subtab: Protein Databases.
9. Press the link: Import.
10. In popup window, select Human protein database.
11. In popup window, select Parsing Rule: Uniprot FASTA.
12. In popup window, press the link: Import.
13. Repeat steps IV – 9, 10, 11, and 12 for Yeast protein database, and then E. Coli protein database.
14. Select the tab: Report.
15. Press the link: Import Schema.
16. In popup window, select protein report template.
17. Press the button: Open.
18. Repeat steps IV – 15, 16, and 17 for peptide report template.

#### V) Data processing of QC sample analyses with Spectronaut

1. Start the program "Spectronaut".
2. Select the tab: Review.
3. Press the link: Load Raw from File
    a. In popup window, select the three QC analyses replicate.
    b. In popup window, press the button: Open
4. In window "Experiment Setup":
    a. Enter as Experiment name "Lab_W-Day_X-Sample_QC" (where "W" is the lab number [*e.g.*, "1" or "2"] and "X" is the day number [1, 3, 5, or 7]).
    b. Select the Tab: FASTA Files
    c. Click in checkbox of Human Protein Database
    d. Press the Link: Configure Conditions
    e. In window "Condition Editors", Enter "Sample_QC" in 3 cells of column "Condition" (each sample QC raw file row). Press the link: Apply
    f. Press the Link: Spectral Library"
    g. In window "Import Spectral Library", select tab "From Prepare Perspective". Select Human Spectral Library. Press link: Load.
    h. Select the Tab: Analysis Settings.
    i. Select Schema "BGS Factory Settings (default") if not already selected.
    j. Verify that all parameters of the different subsections of "BGS Factory Settings" are set at default values (refer to screenshot below, section VIII). Check more specifically that in subsection "Quantification", parameter "Quantify MS-Level" is set at "MS1".
    k. Press the link: Start.
5. At the end of data processing, righ-click in the left panel (tab including experiment name) and select "Save as". In popup window, enter "Lab_W-Day_X-Sample_QC-SpectronautAnalysis" (where "W" is the lab number [*e.g.*, "1" or "2] and "X" is the day number [1, 3, 5, or 7]) and press button: Save.
6. Select Tab: Report
7. In left panel "Schemas", select QuanBenchmark_proteins
8. Press the link: Export Report. In popup window, enter "Lab_W-Day_X-Sample_QC-ReportProteins" (where "W" is the lab number [*e.g.*, "1" or "2"] and "X" is the day number [1, 3, 5, or 7]) and press button: Save.
9. In left panel "Schemas", select QuanBenchmark_peptides
10. Press the link: Export Report. In popup window, enter "Lab_W-Day_X-Sample_QC-ReportPeptides" (where "W" is the lab number [*e.g.*, "1" or "2"] and "X" is the day number [1, 3, 5, or 7]) and press button: Save.

#### VI) On-site evaluation of QC sample analyses with Spectronaut

1. In Spectronaut, open Spectronaut file "Lab_W-Day_X-Sample_QC-SpectronautAnalysis" if not opened yet.
    a. Select the tab: Review
    b. Press the link: Load Spectronaut Experiment.
    c. In popup window, select file "Lab_W-Day_X-Sample_QC-SpectronautAnalysis"
    d. In popup window, press the button: Open.
2. In Spectronaut, select the tab: Review.
    a. In main window, select "Analysis Summary" in drop-down menu "Lower panel".
    b. Extract values indicated in lower panel for "Median Peak Width", "Data Points per Peak (MS1)", and "Data points per Peak (MS2).

3. In Spectronaut, select the tab: Post Analysis.
    a. In left panel, in subsection “Analysis Overview”, select “Coefficients of Variation”. Precursor CV distribution graph is displayed, including median value. By right clicking in the graph and selecting “CV Base” -> “Peptide” or “Protein Group”, Peptide or Protein Group CV graph can be displayed.
    b. Extract the different median values.
4. In Spectronaut, in the tab: Post Analysis / left panel / subsection “Analysis Overview”, select “CVs below X”.
    a. Graph “Precursor CVs below X” is displayed. By right clicking in the graph and selecting “Show Point Values”, direct access to numerical values are provided by moving the cursor over the different columns of the histogram (numbers of precursors “identified” / “identified with CV<20%” / “identified with CV<10%”). By right clicking in the graph and selecting “CV Base” -> “Peptide” or “Protein Group”, graphs “Peptide CVs below X” or “Protein Group CVs below X” can be displayed.
    b. Extract the numbers of precursors/peptides/proteins “identified”, “identified with CV<20%”, and “identified with CV<10%”.

#### VII) Centralized data processing of Samples A and B with Spectronaut

The following procedure was only applied at one lab were all the data were centralized.

1. Start the program “Spectronaut”.
2. Select the tab: Review.
3. Press the link: Load Raw from File
    a. In popup window, select the three Sample A analyses and three Sample B analyses for each lab and day retained.
    b. In popup window, press the button: Open
4. In window “Experiment Setup”:
    a. Enter as Experiment name “MultiSite-Sample_A-B”.
    b. Select the Tab: FASTA Files
    c. Click in checkbox of Human Protein Database, Yeast Protein Database, and E. Coli Protein Database.
    d. Press the Link: Configure Conditions
    e. In window “Condition Editors”, Enter “Lab_W-Day_X-Sample_Y” (where “W” is the lab number [*e.g.*, “1” or “2”], “X” is the day number [1, 3, 5, or 7], and “Y” is the sample name [A or B]) in the pertinent cells of column “Condition”. Press the link: Apply.
    f. Press the Link: Spectral Library”
    g. In window “Import Spectral Library”, select tab “From Prepare Perspective”. Select Human Spectral Library. Press link: Load.
    h. Repeat step VII 4 – g for Yeast Spectral library, and then E. Coli Spectral Library.
    i. Select the Tab: Analysis Settings.
    j. Select Schema “BGS Factory Settings (default”) if not already selected.
    k. Verify that all parameters of the different subsections of “BGS Factory Settings” are set at default values (refer to screenshot below, section VIII). Check more specifically that in subsection “Quantification”, parameter “Quantify MS-Level” is set at “MS1”.
    l. Press the link: Start.
5. At the end of data processing, righ-click in the left panel (tab including experiment name) and select “Save as”. In popup window, enter “MultiSite-Sample_A-B” and press button: Save.
6. Select Tab: Report
7. In left panel “Schemas”, select QuanBenchmark_proteins

8. Press the link: Export Report. In popup window, enter “ID_Controlled-Samples_MultiSite_ReportProteins” and press button: Save.
9. In left panel “Schemas”, select QuanBenchmark_peptides
10. Press the link: Export Report. In popup window, “ID_Controlled-Samples_MultiSite_ReportPeptides” and press button: Save.

#### VIII) Screenshots required values for parameters in Tab “Analysis Settings” / Schema “BGS Factory Settings” / Subsection “Quantifications” (unless specified differently)

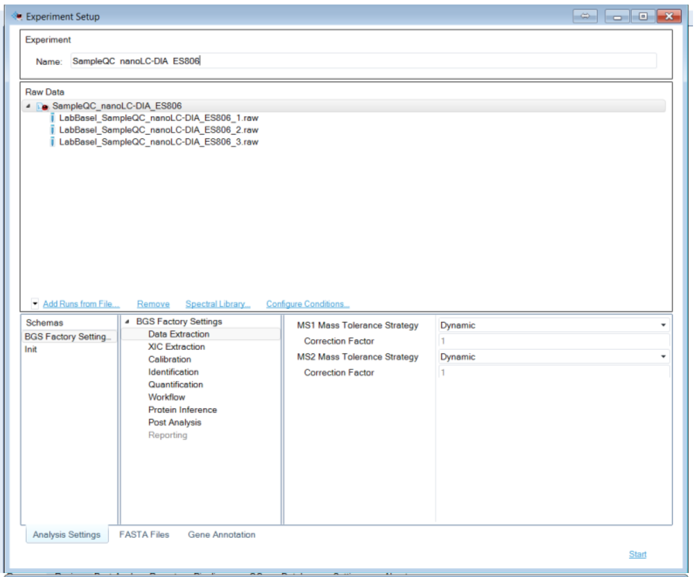


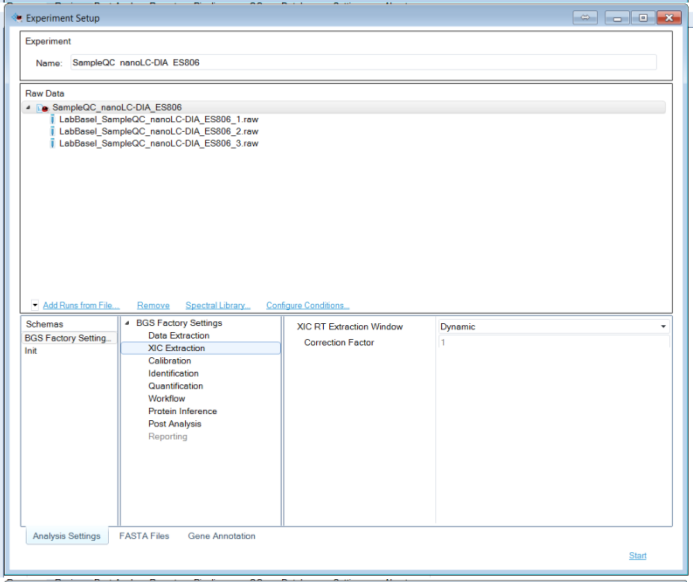


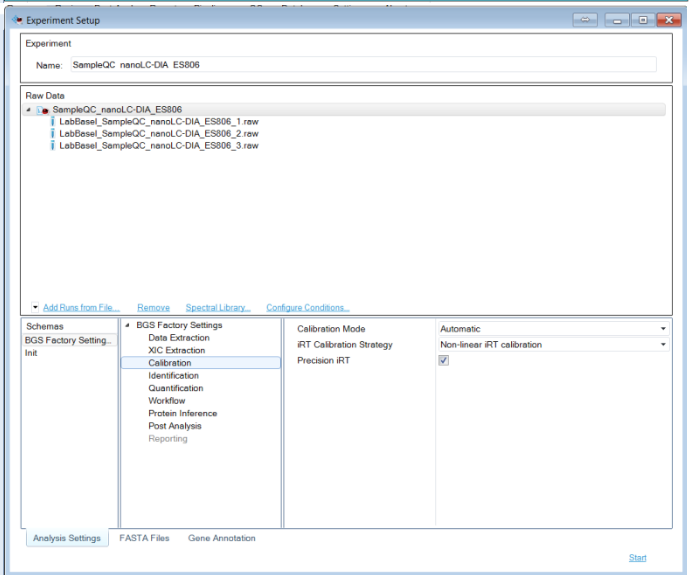

#### Cancer Moonshot Multiple Sites Study

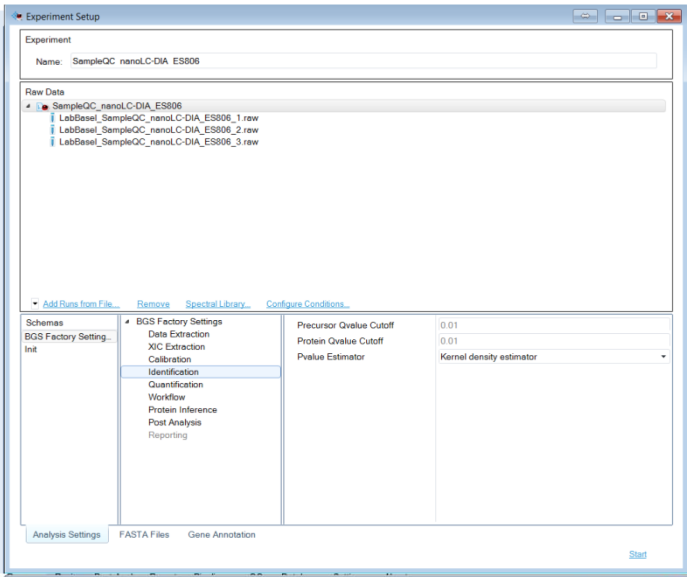


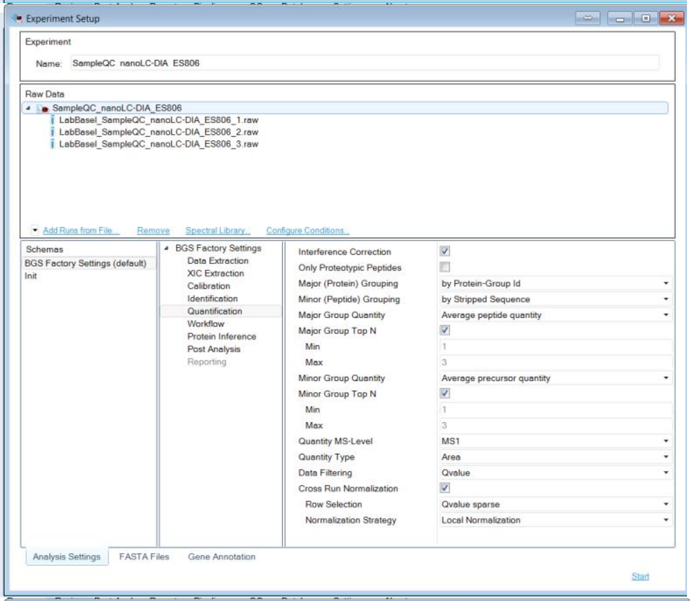


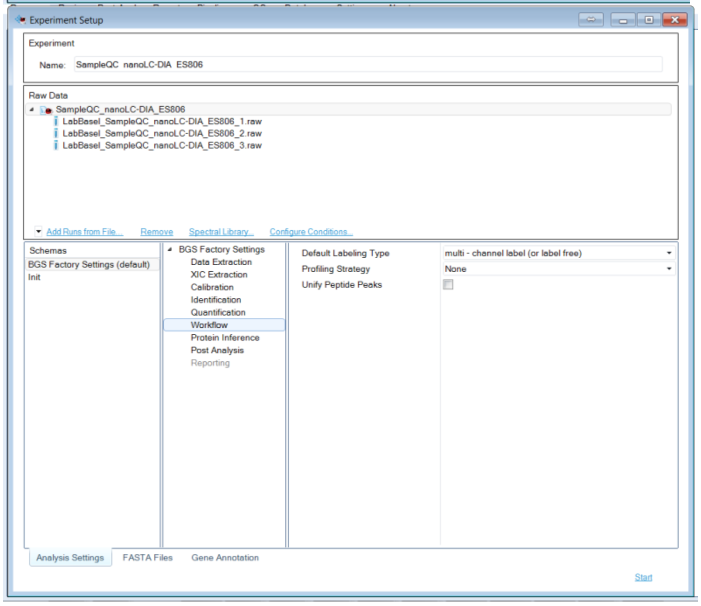

#### Cancer Moonshot Multiple Sites Study

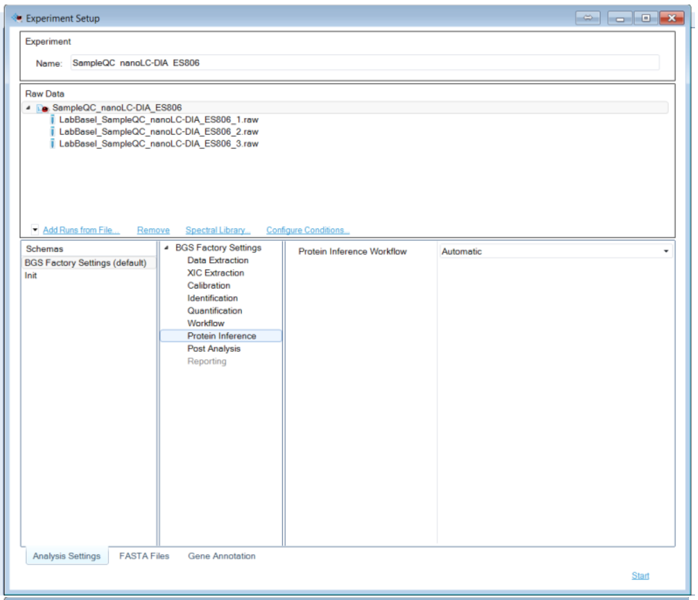


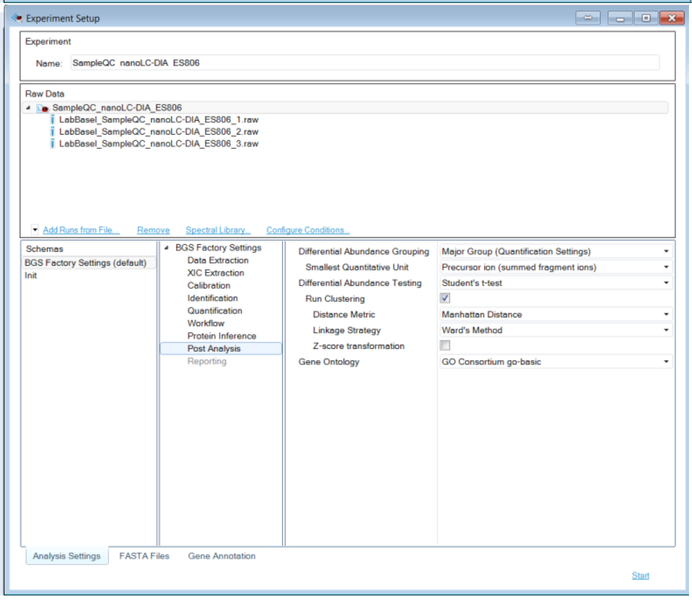
